# Functional Landscape of Motifs within the Sarbecovirus Spike Cytoplasmic Tail

**DOI:** 10.64898/2026.05.06.723231

**Authors:** Shixiong Zhou, Chong Chen, Chunhai Liu, Xinhao Zheng, Lanfang Shi, Lina Ma, Pengfei Cheng, Qian Wang, Lihong Liu

**Author notes:** Correspondence, (L.L.). These authors contributed equally (Shixiong Zhou, Chong Chen).

## Abstract

Sarbecoviruses exhibit extensive diversity in host range and zoonotic potential. Although the ectodomain of the viral spike protein has been well characterized, the functional landscape of the cytoplasmic tail (CT) remains poorly defined. To address this gap, we systematically generated CT truncation variants of representative spike proteins from all major ACE2-utilizing clades to define the roles of conserved host factor–interacting motifs associated with COPI, COPII, FERM, and SNX27. Using vesicular stomatitis virus (VSV)- and lentivirus-based pseudotyping systems, we evaluated viral entry in cells expressing varying levels of ACE2 and TMPRSS2. Our results demonstrate that CT-associated motifs differentially regulate viral infectivity. Specifically, truncation of the COPI- or SNX27-binding motifs markedly reduces entry efficiency, whereas disruption of the COPII-binding motif produces the opposite outcome. By contrast, removal of the FERM-binding motif consistently enhances infectivity across lineages. Mechanistically, truncation of this motif increases spike expression, cell surface localization, incorporation into virions, and particle stability. Importantly, despite these pronounced effects on viral infectivity, deletion of the FERM-binding motif does not affect antigenicity, receptor dependence, or sensitivity to protease inhibitors, as demonstrated by neutralization and inhibition assays. In addition, this approach substantially increases spike protein density on virus-like particles (VLPs). Collectively, by extending the analysis beyond SARS-CoV-1 and SARS-CoV-2, our study reveals a generalizable mechanism in which cytoskeletal anchoring mediated by the FERM-binding motif acts as a limiting determinant of viral assembly. These findings provide a practical framework for optimizing pseudovirus platforms and guiding vaccine development against emerging viral threats.

## Introduction

Sarbecoviruses, a subgenus within lineage B of the genus *Betacoronavirus,* are predominantly hosted by bats and other natural reservoirs and possess zoonotic spillover potential with the capacity to infect humans^1, 2, 3, 4, 5, 6^. Over the past two decades, two members of this group, severe acute respiratory syndrome coronavirus 1 and 2 (SARS-CoV-1 and SARS-CoV-2), have caused an epidemic and a pandemic, respectively, underscoring the substantial threat posed by this subgenus to public health worldwide^7, 8^. In this context, elucidating the molecular determinants underlying viral transmission and host infection remains essential. The viral spike protein is central to these processes.

Structurally, the coronavirus spike protein is a typical type I transmembrane protein that forms homotrimers and comprises a large extracellular domain at the N-terminus, a transmembrane domain, and a short cytoplasmic tail (CT) at the C-terminus^9, 10, 11, 12, 13,14^. The ectodomain, consisting of the S1 subunit and the majority of the S2 subunit, is responsible for receptor binding and membrane fusion, and has therefore been extensively characterized as a principal target for vaccines and antiviral therapeutics^15, 16, 17, 18^. Phylogenetic analysis of nucleotide sequences encoding the receptor-binding domain (RBD) of the spike protein indicates that sarbecoviruses can be classified into four major groups (Clades 1-4) (**Fig. 1a**), which exhibit substantial divergence in receptor usage and host tropism. Notably, viruses from Clades 1, 3, and 4 have been shown to utilize angiotensin-converting enzyme 2 (ACE2) from a wide range of host species for cell entry. In contrast, the receptor usage and potential for cross-species transmission of Clade 2 viruses remain poorly defined and warrant further investigation^1, 19, 20, 21^. However, the functional properties of the CT are relatively underexplored. Existing studies indicate that the CT of the spike protein harbors multiple of linear motifs (**Fig. 1b**) that interact with host proteins such as COPI (coat protein complex I), COPII (coat protein complex II), FERM (4.1 protein, ezrin, radixin, and moesin) domain-containing proteins, and SNX27 (sorting nexin 27)^22, 23, 24, 25, 26, 27,28^. These motifs collectively are involved in intracellular trafficking of the spike protein, as well as in viral assembly and budding.

**Fig. 1.**
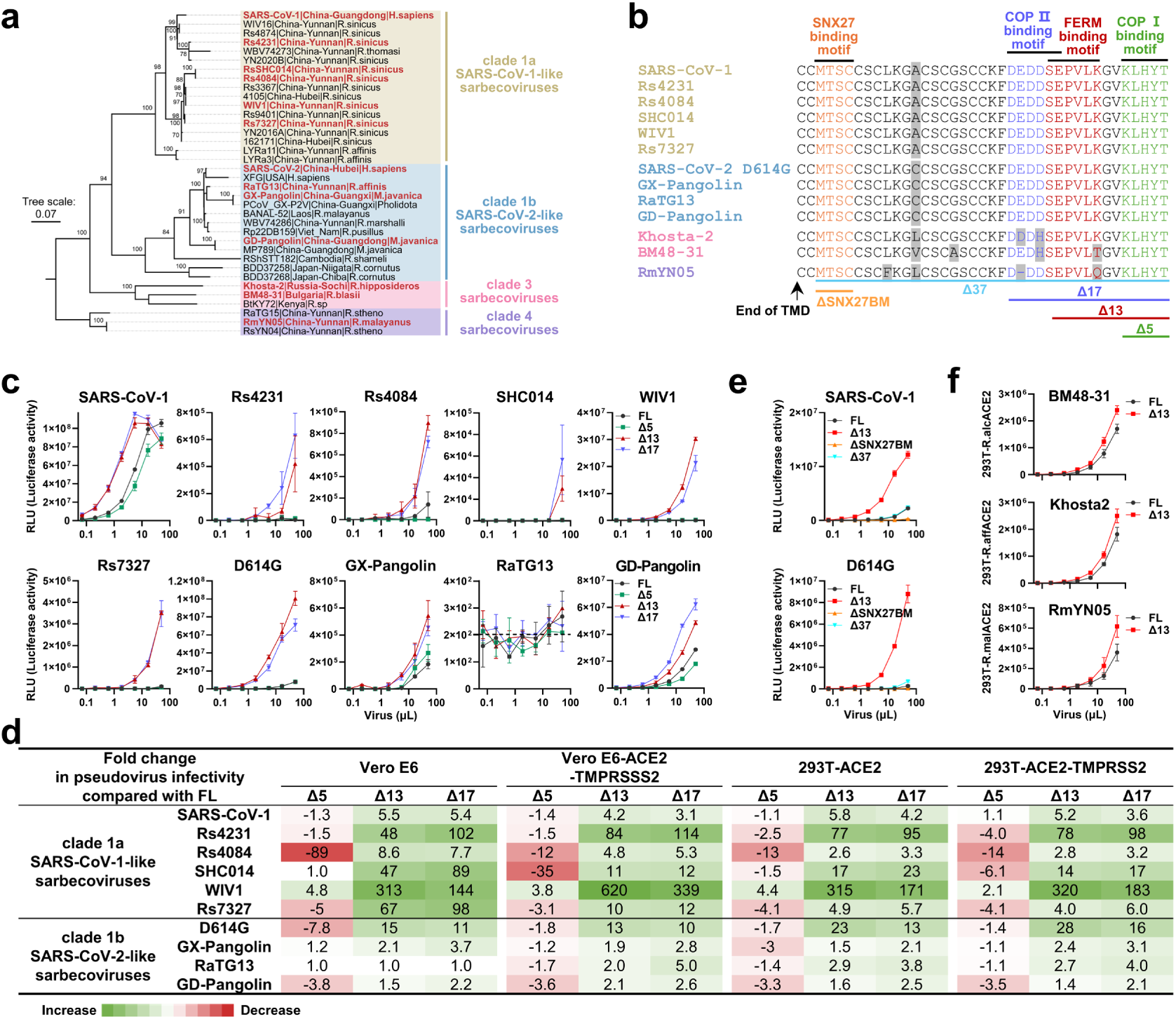
Cytoplasmic tail (CT) truncations of spike glycoproteins increase VSV pseudovirus titers of sarbecoviruses. **a.** Maximum-likelihood phylogenetic tree of representative sarbecoviruses, including clade 1a (SARS-CoV-1-like), clade 1b (SARS-CoV-2-like), clade 3, and clade 4 viruses. The tree was generated based on amino acid sequences of the spike RBD. Strains selected as representatives in this study are highlighted in red. Tip labels are formatted as ‘strain name | discovery location | host’. **b.** CT sequence alignment of the indicated spikes. Solid black lines annotate COPI, COPII, and FERM-binding motifs. Deletion of 5, 13, and 17 residues (Δ5, Δ13, and Δ17) at the C-terminus of the CT are also indicated. TMD, transmembrane domain. Δ, deletion; SNX27, sorting nexin 27; COPI, coat protein complex I; COPII, coat protein complex II; FERM, 4.1/Ezrin/Radixin/Moesin; ΔSNX27BM, deletion of SNX27-binding motif. **c.** Titration of VSV pseudoviruses carrying FL or truncated spike proteins (Δ5, Δ13, and Δ17) in Vero-E6 cells. Infectivity was assessed by measuring luciferase activity (relative light units, RLU). Δ13 and Δ17 pseudoviruses, both lacking a FERM-binding motif, exhibited higher infectivity than FL and Δ5 pseudoviruses, indicating a key role for the FERM-binding motif in regulating infectivity. **d.** Fold changes in pseudovirus infectivity of VSV pseudotyped with Δ5, Δ13, and Δ17 spike compared with that of the FL spike in panel **c**. Fold changes were derived from the linear range of infectivity values. Increased infectivity is in green and decreased in red. **e.** Titration of VSV pseudoviruses of SARS-CoV-1 and SARS-CoV-2 D614G carrying FL or truncated spike proteins (Δ5, Δ13, ΔSNX27BM, and Δ17) in Vero E6 cells. **f.** Titration of VSV pseudoviruses of BM48-31, Khosta-2, and RmYN05 carrying FL or Δ13 spike in 293T cells transiently expressing ACE2 of *R. alcyone* (R.alcACE2), *R. affinis* (R.affACE2), and *R. malayanus* (R.malACE2).

In the classical vesicular transport pathway, COPI- and COPII-binding motifs within the spike protein CT coordinately regulate its intracellular trafficking. The COPI-binding motif mediates retrograde transport from the Golgi apparatus to the endoplasmic reticulum (ER), thereby promoting spike accumulation at the viral assembly site, the ER-Golgi intermediate compartment (ERGIC)^29, 30, 31^. In sarbecoviruses, the motif has evolved into a noncanonical KxHxx configuration (e.g., “KLHYT”), whereas canonical COPI-binding motifs typically comprise dibasic sequences, including KxKxx or KKxx^32, 33^. Specifically, within noncanonical motif, the second functionally relevant residue is substituted from lysine (K) to histidine (H), which reduces its COPI-mediated ER retrieval efficiency mediated by COPI complex proteins^22, 34, 35, 36^. Such attenuated retrieval appears to favor viral assembly, as observed in SARS-CoV-2, whereas reverting histidine to lysine enhances ER retention of spike and impairs its glycosylation maturation and proteolytic processing^37^. In contrast to COPI-mediated retrograde transport, the COPII-binding motif is primarily involved in the anterograde secretory pathway of spike. The CT of spike contains a typical COPII-binding motif characterized by an acidic di-acidic sequence, commonly with the consensus DxE motif^38^. In SARS-CoV-2, this region manifests as “DEDDSE,” which binds to the Sec24 subunit of the COPII complex, thereby mediating ER-to-Golgi export of spike and enabling subsequent proteolytic cleavage and glycosylation^22^. Together, COPI- and COPII-mediated bidirectional trafficking pathways ensure the proper distribution and maturation of the spike protein within the secretory system.

Beyond canonical vesicular trafficking signals, the spike CT also contains a FERM-binding region capable of interacting with host ERM (ezrin/radixin/moesin) family^39^. ERM proteins bind transmembrane proteins via their N-terminal FERM domain and link them to F-actin (filamentous actin) through their C terminus, thereby anchoring membrane proteins to the cytoskeleton, regulating their stability and membrane distribution, and participating in signal transduction^40, 41^. This binding site does not possess a strictly conserved short linear consensus sequence. In SARS-CoV-2, the interaction involves the “SEPVLK” sequence located in the juxtamembrane region of spike protein. Li et al. identified that P1263 is a determinant residue for FERM binding, and its substitution with leucine (L) markedly weakens the interaction between spike and ERM proteins^27^. This suggests that the region plays a vital role in coupling membrane proteins to the cytoskeletal framework.

In addition, the CT of sarbecovirus spike proteins possesses a noncanonical SNX27-binding motif. Although it does not fully conform to the typical PSD95, Dlg1, zo-1 (PDZ)-binding motif consensus sequence (X-S/T-X-Φ, where Φ denotes a hydrophobic residue), in vitro GST pull-down assays have confirmed its interaction with the host protein SNX27^25, 26, 42, 43^. SNX27, a member of the sorting nexin family, recognizes PDZ-binding motifs in the cytosolic regions of target proteins via its PDZ domain, thereby mediating post-endocytic recycling and trafficking proteins from endosomes to the plasma membrane or the trans-Golgi network^43^. It is worth noting that the SNX27- binding motif is located in the C-terminal region of the CT, proximal to the transmembrane domain, which is enriched in cysteine residues and represents a hotspot for palmitoylation. The lipid modifications may confer membrane-anchoring properties to this region and influence the spatial accessibility of the SNX27-binding site^22, 44^. Consequently, the physiological function and regulatory mechanisms of the SNX27-binding motif within the cellular environment of sarbecoviruses have yet to be determined.

The aforementioned studies indicate that the CT is a key regulator of viral assembly in SARS-CoV-2. Truncation of specific regions within the CT can also considerably increase the titers of SARS-CoV-1 and SARS-CoV-2 pseudoviruses and enhance spike incorporation into virus-like particles (VLPs)^45, 46, 47, 48^. However, research on various motifs within the spike CT continues to be mainly centered on SARS-CoV-2. Systematic comparisons evaluating the effects of these motifs on viral infectivity and particle stability across different sarbecovirus lineages are still lacking. This knowledge gap hinders a comprehensive understanding of the general regulatory principles and evolutionary features of the spike CT in sarbecoviruses.

To address these questions, we selected representative spike proteins from distinct sarbecovirus clades based on phylogenetic analysis, generated CT truncation mutants, and systematically compared the biological properties of wild-type and truncated variants using both lentiviral- and VSV-based pseudoviruses. Our results demonstrate that removal of the FERM-binding motif, instead of the COPI-, COPII-, or SNX27-binding motifs, significantly enhances spike protein expression, viral infectivity, and viral particle stability without compromising antigenicity. Consistently, truncation of the FERM-binding motif robustly promotes spike incorporation into VLPs, highlighting its regulatory activity in spike trafficking and virion assembly. Overall, these findings extend the mechanistic insight of spike CTs beyond SARS-CoV-2, revealing conserved functions in viral particle assembly across sarbecoviruses. Importantly, this work establishes a practical for optimizing pseudovirus platforms, enabling robust in vitro infection models for entry assays and neutralization studies, particularly for low-infectivity viruses, and advancing vaccine design for emerging viral threats and pandemic preparedness.

## Results

### CT truncation of sarbecovirus spike glycoproteins enhances VSV pseudovirus titers

Previous studies have shown that truncation of the CT of the SARS-CoV-2 spike can significantly enhance viral titers^47^. Both SARS-CoV-1 and SARS-CoV-2 are classified within the subgenus sarbecovirus of the *Coronaviridae* family (**Fig. 1a**). In this study, we sought to determine whether CT truncation-mediated enhancement of viral titers can be extended to other sarbecoviruses beyond SARS-CoV-2. Accordingly, we selected six spike proteins from clade 1a and four from clade 1b, together with three additional spikes from clades 3 and 4. The representative spike proteins analyzed in this study are highlighted in red in the phylogenetic tree (**Fig. 1a**). Clade 2 spikes were excluded because their receptor remains unknown and no suitable target cells are currently available for viral entry^49^. Amino acid sequence alignment revealed that CT sequences are relatively conserved among the selected sarbecoviruses, with only a single amino acid difference between clade 1a and clade 1b spikes (**Fig. 1b**), and 3-4 residue differences between clade 1 and clades 3/4 spikes.

The CTs of sarbecovirus spikes have been reported to contain SNX27-, COPI-, COPII-, and FERM-binding motifs (**Fig. 1b**). To examine the impact of these motifs on viral infectivity, we generated C-terminal truncations of 5, 13, 17, and 37 amino acids (Δ5, Δ13, Δ17, and Δ37), thereby removing one or more motifs. Full-length (FL) and truncated spikes were subsequently incorporated into VSV pseudotyped viruses. The corresponding pseudoviruses were then titrated on Vero E6 cells, a widely used cell line in SARS-CoV-2 research^45, 50, 51, 52^ and infectivity was quantified by measuring luciferase activity (**Fig. 1c**). Pseudoviruses carrying the FL spikes of Rs4231, SHC014, and RaTG13 exhibited nearly undetectable viral infection in Vero E6 cells with the maximal readouts of luciferase activity (relative light units, RLU) ranging from 268 to 4,927 (**Fig. 1c**), thereby limiting the evaluation of their infectivity and neutralization sensitivity.

To enhance target cell susceptibility, we transduced Vero E6 and 293T cells with the clade 1 sarbecovirus receptor ACE2, either alone or in combination with the host protease TMPRSS2 (transmembrane protease serine 2), which promotes S2′ cleavage and facilitates viral entry^53^. The resulting Vero E6-ACE2-TMPRSS2, 293T-ACE2, and 293T-ACE2-TMPRSS2 cell lines were evaluated for ACE2 and TMPRSS2 expression by flow cytometry **(Fig. S1a,b)**. Viruses were then titrated on these three cell lines and viral infectivity is shown in **Fig. S1c-e**. The presence of high levels of ACE2 and/or TMPRSS2 increased viral infectivity by 1.1- to 3,043-fold in Vero E6 cells and 3.0- to 122-fold on 293T-ACE2 cells (**Fig. S1f**). Notably, all ten pseudoviruses carrying FL spikes exhibited measurable luciferase activity (RLU >10^5^ and >100-fold above the background) on the three engineered cell lines (**Fig. S1g**).

The infectivity of pseudoviruses bearing truncated spikes in Vero E6, Vero E6-ACE2-TMPRSS2, 293T-ACE2, and 293T-ACE2-TMPRSS2 cells was normalized to that of their corresponding FL counterparts (**Fig. 1d**). The Δ13 and Δ17 truncations enhanced pseudovirus infectivity by 1.4- to 620-fold across all four cell lines, except for RaTG13 on Vero E6 (**Fig. 1d**). Although the Δ13 and Δ17 variants of RaTG13 did not increase infectivity in Vero E6 cells, their titers improved by 2.0- to 5.0-fold on the engineered Vero E6-ACE2-TMPRSS2, 293T-ACE2, and 293T-ACE2-TMPRSS2 cells (**Fig. 1d**).

In contrast to Δ13 and Δ17, both of which remove the FERM-binding motif, Δ5 consistently reduced pseudovirus titers, with WIV1 being the only exception. These findings suggest that deletion of the COPI-binding motif (Δ5) alone impairs infectivity, whereas the removal of the terminal 13 or 17 residues (Δ13 and Δ17) significantly enhances it. However, Δ37, which deletes the entire CT, showed pseudovirus infectivity comparable to the FL spike, even though this deletion also contains the FERM-binding motif (**Fig. 1e**). This effect may be attributable to the additional removal of the SNX27-binding motif (ΔSNX27BM) in Δ37, which exerts effects comparable to those of COPI-binding motif deletion (Δ5) (**Fig. 1e**).

In addition to clade 1 viruses, we evaluated the titers of clade 3 and clade 4 viruses (BM48-31, Khosta-2, and RmYN05). Because these bat-origin viruses preferentially utilize their respective host ACE2 receptors, we transiently transfected 293T cells with ACE2 orthologs from *R. alcyone*^54^, *R. affinis*^21^, and *R. malayanus*^55^ to assess FL and Δ13 constructs. Data in **Fig. 1f** show that Δ13 slightly elevated the infectivity of all three viruses compared to their FL counterparts, consistent with the effects observed in clade 1.

To validate these findings, we generated lentiviral pseudotypes for the ten selected spikes in the FL, Δ5, Δ13, and Δ17 contexts. Because Vero E6-derived cell lines express host restriction factors that limit lentiviral infection^56^, lentiviral pseudotypes of SARS-CoV-1 (**Fig. S2a**) and SARS-CoV-2 D614G (**Fig. S2b**) exhibited substantially lower infectivity in Vero E6 and Vero E6-ACE2-TMPRSS2 cells than VSV pseudotypes (**Figs. 1c and S1**). Consequently, the remaining lentiviral pseudotypes were titrated only on 293T-ACE2 (**Fig. S2c**) and 293T-ACE2-TMPRSS2 (**Fig. S2d**) cells. Consistent with results obtained using VSV pseudoviruses, Δ13 and Δ17 lentiviral pseudoviruses yielded higher titers than FL and Δ5 across all ten tested clade 1 sarbecoviruses.

### Enhancement of VSV pseudovirus titers is attributed to removal of the FERM-binding motif

Because Δ13 removes not only the FERM- and COPI- binding motifs but also the last two residues of the COPII-binding motif, it is difficult to determine whether the enhanced viral titers of Δ13 and Δ17 arise from loss of the COPII-binding motif, the FERM-binding motif, or both. To address this, we generated two additional truncations: Δ11, which deletes 11 residues from the CT to disrupt the FERM-binding motif while preserving an intact COPII-binding motif; and ΔFBM, which specifically removes only the FERM-binding motif (**Fig. 2a)**. We then compared their infectivity with that of FL, Δ5, Δ13, and Δ17 spikes in the context of VSV pseudoviruses. As shown in **Figs. 2b,c**, Δ11 exhibited 2.8- to 12-fold higher infectivity than FL for SARS-CoV-1 and SARS-CoV-2 D614G on the four tested cell lines, though its infectivity remained slightly lower (<2-fold difference) than that of Δ13 and Δ17. VSV pseudoviruses carrying the FERM-binding motif deletion (ΔFBM) exhibited a similar pattern in Vero E6 and Vero E6-ACE2-TMPRSS2 cells for SARS-CoV-1, Rs7327, D614G, and GD-pangolin (**Figs. 2d,e**). These data suggest that disruption of the COPII-binding motif in the spike CT slightly promotes infectivity, whereas the deletion of the FERM-binding motif was the primary determinant of increased pseudovirus titers.

**Fig. 2.**
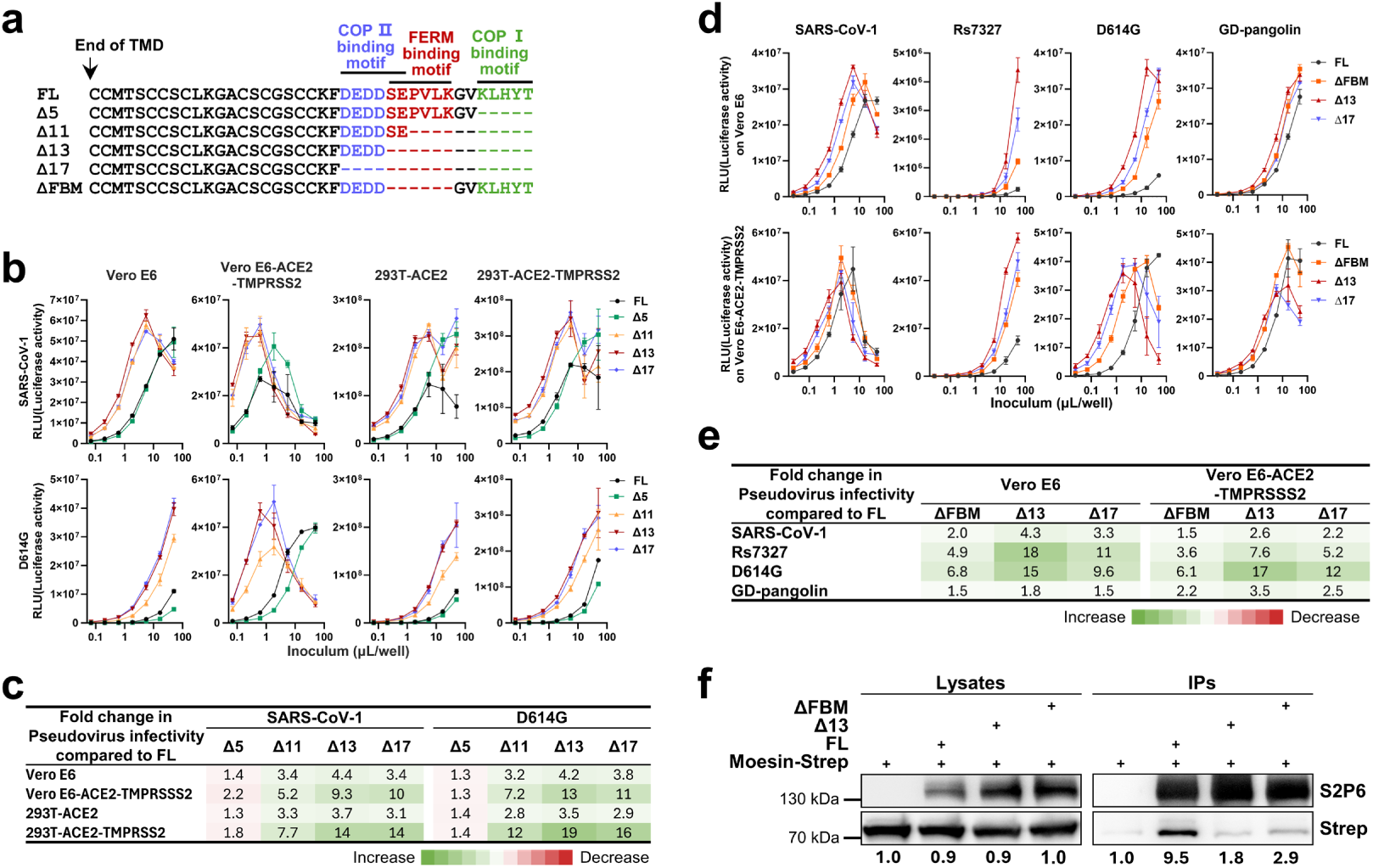
Enhancement of VSV pseudovirus titers is attributed to the FERM-binding motif removal. **a.** Construction scheme of Δ5, Δ11, Δ13, Δ17, and ΔFBM. ΔFBM, deletion of FERM-binding motif. Solid black lines indicate COPI-, COPII-, and FERM-binding motifs. **b.** Titration of VSV pseudoviruses carrying FL or truncated spike proteins (Δ5, Δ13, and Δ17) of SARS-CoV-1 and SARS-CoV-2 D614G in Vero E6 cells. **c.** Fold changes in pseudovirus infectivity of VSV pseudotyped with Δ5, Δ13, and Δ17 spikes compared with that of the FL spike in panel **b**. Fold changes were derived from the linear range of infectivity values. Increased infectivity is shown in green and decreased infectivity in red. **d.** Titration of VSV pseudoviruses carrying FL or truncated spikes (ΔFBM, Δ13, and Δ17) of the indicated viruses in Vero-E6 and Vero E6-ACE2-TMPRSS2 cells. **e.** Fold changes in pseudovirus infectivity of VSV pseudotyped with ΔFBM, Δ13, and Δ17 spike compared with that of the FL spike in panel **d**. Fold changes were derived from the linear range of infectivity values. **f.** Co-immunoprecipitation assays assessing the interaction between moesin and FL or truncated D614G spike proteins. 293T cells were transfected with the indicated plasmids, and cell lysates were subjected to immunoprecipitation using an anti-spike monoclonal antibody (SA55). Protein expression and immunoprecipitation were detected by western blot using an anti-sarbecovirus spike monoclonal antibody (S2P6) and an anti-Strep tag antibody, respectively. Band intensities were quantified using ImageJ software, with both numerical values and color coding representing relative grayscale intensity.

To validate the association between the FERM-binding motif in the spike CT and host ERM family proteins (ezrin, radixin, and moesin), we performed a co-immunoprecipitation (Co-IP) assay in 293T cells (**Fig. 2f**). Briefly, 293T cells were co-transfected with plasmids encoding moesin together with SARS-CoV-2 D614G spike proteins in the FL, ΔFBM, or Δ13 forms, and co-immunoprecipitation of moesin by the respective spike variants was assessed.The results showed that the FL spike efficiently co-immunoprecipitated moesin, whereas deletion of either 13 amino acids from the CT or the FERM-binding motif severely disrupted the interaction. These data indicate that ΔFBM and Δ13 disrupt binding to ERM proteins, thereby promoting spike incorporation into pseudotyped virions and virus-like particles, ultimately enhancing viral infectivity.

### Deletion of the FERM-binding motif in the spike CT facilitates spike expression and pseudoviruses stability

We further investigated the mechanisms by which Δ13 and Δ17 increase pseudotyped sarbecovirus titers while Δ5 has the opposite effect. We assessed spike expression in spike-expressing 293T cells (**Fig. 3a**) and on the surface of 293F cells (**Fig. 3b,c**) for four representative viruses (SARS-CoV-1, Rs7327, D614G, and GD-pangolin). Plasmids encoding FL spikes and the corresponding truncated variants were transiently transfected into 293T cells, and expression was analyzed by western blot using a cocktail of anti-S2 antibodies (S2P6^57^, CV3-25^58^, and CC40.8^59^). Δ13 and Δ17 spikes showed higher expression levels than the FL spike for all four viruses. In contrast, Δ5 spikes exhibited lower expression, with the exception of D614G. However, the cleavage efficiency of the D614G Δ5 spike was markedly lower than that of Δ13 and Δ17, as evidenced by the increased abundance of S2 in the Δ13 and Δ17 samples.

**Fig. 3.**
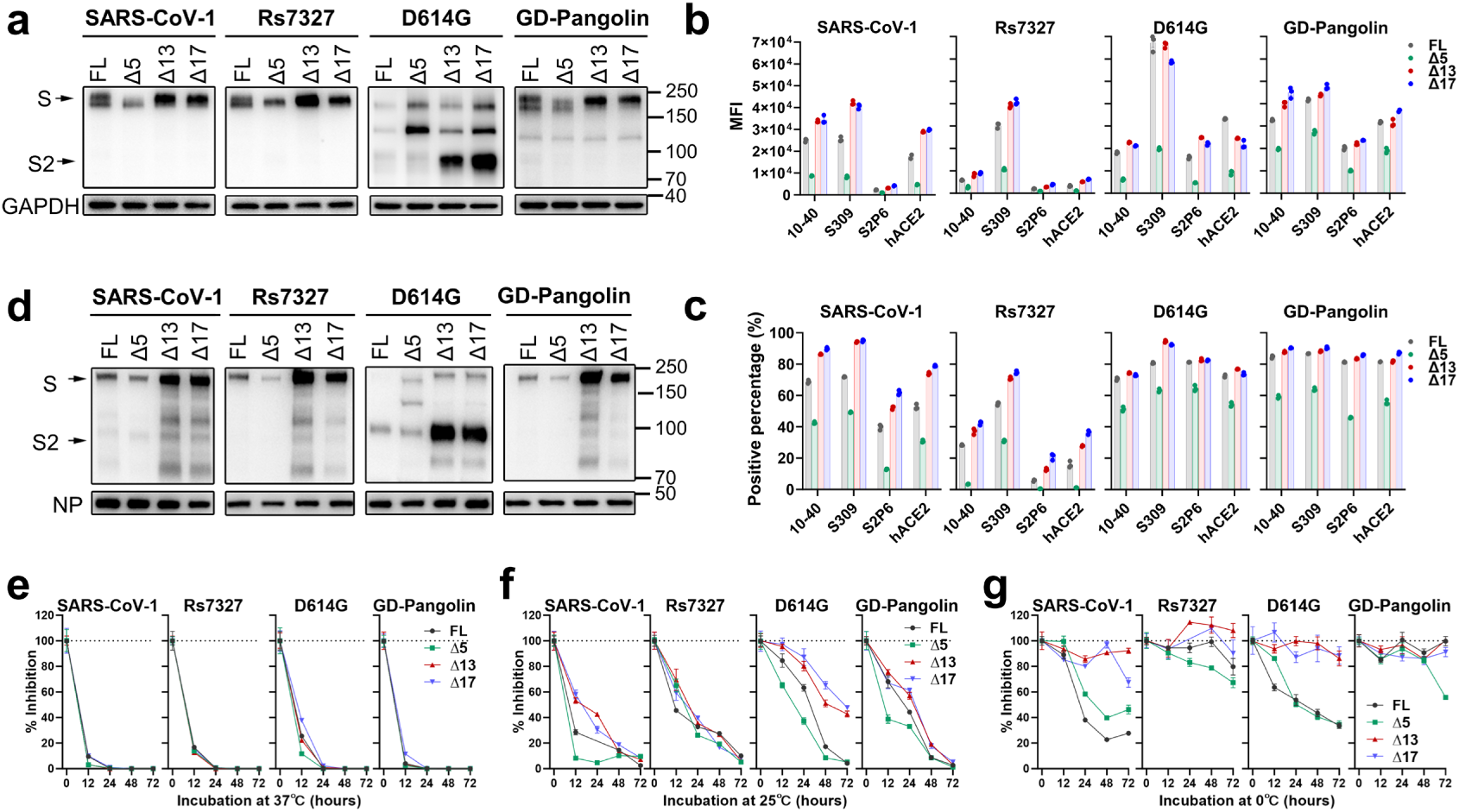
Effects of spike CT truncations on spike expression and stability. **a.** Spike expression in 293T cells transiently transfected with plasmids encoding FL, Δ5, Δ13, and Δ17 spikes. 293T cell lysates were analyzed by western blot using an anti-S2 antibody cocktail (S2P6, CV3-25, and CC40.8). GAPDH served as an internal control. Data shown are representative of three independent experiments. **b-c**. Spike expression on Expi293F cells transiently transfected with plasmids encoding FL, Δ5, Δ13, and Δ17 spikes. Expression of the cell surface spike was assessed by binding of the indicated antibodies and human ACE2 (hACE2) at 10 µg/mL. Data are presented as mean fluorescence intensity (MFI) (**b**) and percentage of positive cells (**c**) from two technical replicates and are representative of two independent experiments. **d.** Incorporation of spike into VSV particles. Samples were analyzed by western blot and probed with anti-S2 antibody cocktails (S2P6, CV3-25, and CC40.8), as well as anti-VSV nucleocapsid protein polyclonal antibodies. Data shown are representative of three independent experiments. **e-g.** Stability of FL, Δ5, Δ13, and Δ17 spikes at 37 °C (**e**), 25 °C (**f**), and 0 °C (**g**). VSV pseudotypes bearing the indicated spike proteins were incubated at the specified temperatures for various time points. Spike stability was calculated based on the infectivity of the pseudoviruses on Vero E6 cells. Relative infectivity normalized to time 0 was calculated and is presented as means ± s.e.m. of three replicates. Results are representative of two independent experiments.

We then quantified spike protein expression on the surface of transiently transfected 293F cells by assessing binding to spike-specific antibodies targeting various epitopes, as well as to human ACE2 (hACE2), using flow cytometry. Antibodies 10-40^60^ and S309^61^ are directed to class 1 and class 3 epitopes of the spike RBD, respectively, and S2P6^57^ targets a linear epitope in S2. Binding was reported as mean fluorescence intensity (MFI) in **Fig. 3b and Fig. S3a,** and percentage of positive cells in **Fig. 3c and Fig. S3b**. The spike expression pattern on the cell surface was overall consistent with the 293T cellular expression data. Δ5 spikes showed lower MFI and positivity for 10-40, S309, S2P6, and hACE2 binding than FL spikes for all four viruses tested, indicating that the cell surface expressing of Δ5 spikes is lower. In contrast, Δ13 and Δ17 spikes exhibited comparable or slightly higher binding to the tested antibodies and hACE2 than FL spikes.

We also measured spike incorporation into VSV pseudotypes via western blot, with data presented in **Fig. 3d**. VSV nucleocapsid protein (NP) was used as an internal control. Notably, Δ13 and Δ17 spike proteins were more efficiently incorporated into virions than FL and Δ5 spike proteins, as indicated by the higher intensity of S or S2 bands. The increased incorporation of Δ13 and Δ17 spike proteins into VSV pseudoviruses correlates with their enhanced infectivity across multiple cell lines. Additionally, Δ5 exhibited slightly reduced spike incorporation relative to FL, which may account for the lower infectivity of Δ5 pseudoviruses.

In addition to assessing spike expression and incorporation, we evaluated spike stability at 37 °C (**Fig. 3e)**, 25 °C (**Fig. 3f**), and 0 °C (**Fig. 3g**) over various time periods in the context of pseudoviruses by measuring relative infectivity after incubation. All four spike variants of SARS-CoV-1, Rs7327, D614G, and GD-pangolin were less stable at 37 °C than at 25 °C and 0 °C, with all variants showing marked instability after 24 h of incubation. At 25 °C, Δ5 pseudoviruses of SARS-CoV-1, D614G, and GD-pangolin were less stable than their FL counterparts, whereas Δ13 and Δ17 were more stable. No major differences in stability were observed among FL, Δ5, Δ13, and Δ17 pseudoviruses of Rs7327. After incubation at 0 °C, Δ13 and Δ17 spikes were generally more stable than the FL and Δ5 spikes. In summary, deletion of the FERM-binding motif in the spike C-terminal tail (Δ13 and Δ17) increased spike expression, enhanced incorporation into virions, improved spike stability, and resulted in higher infectivity of VSV pseudoviruses.

### FERM-binding motif deletion does not alter spike antigenicity or protease inhibitor susceptibility

Data presented in **Figs. 1-3** suggest that deletion of the FERM-binding motif increases viral titers by enhancing spike expression, promoting incorporation into virions, and stabilizing the spike protein. Next, we aimed to determine whether such deletion affects the antigenicity or protease inhibitor susceptibility of sarbecoviruses. To this end, we selected four representative pseudoviruses with high titers in the FL, Δ13, and Δ17 forms and tested their neutralization sensitivity to a panel of five antibodies targeting the spike RBD (**Fig. 4a**), as well as seven serum samples collected after the COVID-19 pandemic (**Fig. 4b**), with clinical information summarized in **Table S1**.

**Fig. 4.**
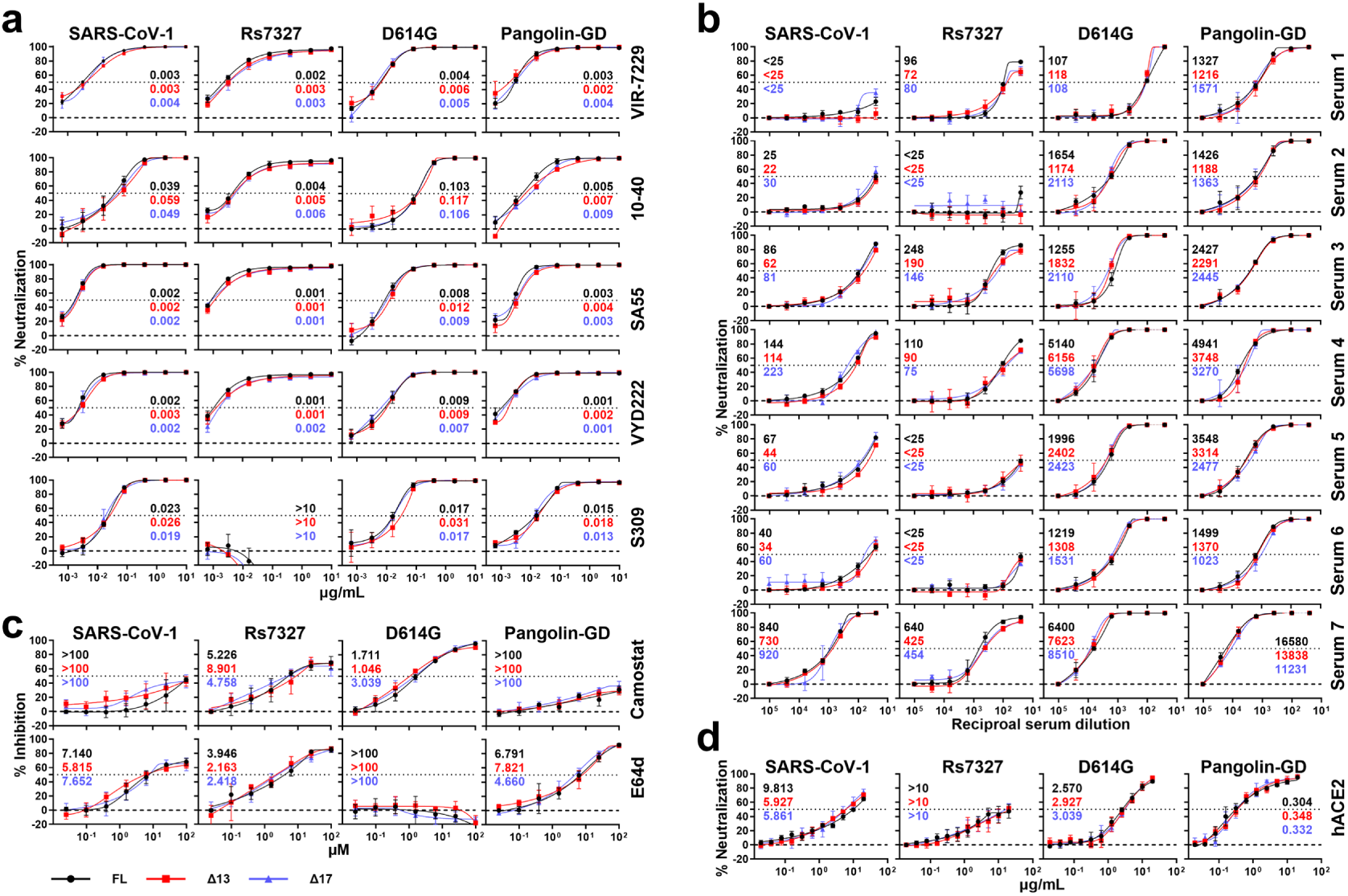
Δ13 and Δ17 spikes exhibit comparable antigenicity and protease inhibitor susceptibility to FL spike. Neutralization curves of VSV pseudoviruses carrying FL, Δ13, and Δ17 spike to the indicated monoclonal neutralizing antibodies (**a**), serum samples collected after SARS-CoV-2 pandemic (**b**), hACE2 (**c**), and the protease inhibitors E64d and camostat (**d**). Antibody and serum neutralization assays, as well as hACE2 inhibition assays, were performed in Vero E6 cells, whereas protease inhibitor assays were conducted in Vero E6-ACE2-TMPRSS2 cells. Numbers in each panel denote IC_50_ or ID_50_ of pseudoviruses carrying FL (black), Δ13 (red), and Δ17 (blue) spikes.

The antibody panel included VIR-7229^62^, 10-40^60^, SA55^63^, VYD222^64, 65^, and S309^61^, all of which have been reported to effectively neutralize both SARS-CoV-1 and SARS-CoV-2. However, S309 failed to neutralize Rs7327 with a 50% inhibitory concentration (IC_50_) greater than 10 µg/ml. As expected, all four viruses showed similar IC_50_ values to antibody neutralization across the FL, Δ13, and Δ17 formats, with less than a 2-fold difference among them. In serum neutralization assays, both clade 1b viruses (D614G and GD-Pangolin) showed higher susceptibility than SARS-CoV-1 and Rs7327, likely because all sera were obtained from donors vaccinated against or previously infected with SARS-CoV-2, and GD-pangolin is more genetically close to SARS-CoV-2. Although some differences in serum ID_50_ titers were observed between the FL, Δ13, and Δ17 formats, these differences were not significant, with fold changes of less than 2-fold as well.

We also evaluated the susceptibility of the FL, Δ13, and Δ17 viruses to hACE2 - mediated (**Fig. 4c**) and protease-mediated (**Fig. 4d**) inhibition using SARS-CoV-1, Rs7327, D614G, and GD-pangolin. Both Δ13 and Δ17, which lack FERM-binding motif, did not alter receptor-mediated inhibition across the four viruses tested (**Fig. 4c**). Upon binding to human ACE2, the second protease cleavage site (S2’) is exposed and recognized by either TMPRSS2 on the cell surface or cathepsin L in the endosome, or both^66^. The protease activity of TMPRSS2 and cathepsin L can be inhibited by camostat^67^ and E64d^68^, respectively. Compared with the FL, Δ13 and Δ17 showed similar sensitivity to Camostat and E64d inhibition, indicating that after the conformational change induced by ACE2 binding, the S2’ protease cleavage site was exposed to a similar degree in FL, Δ13, and Δ17 spike proteins.

Data from monoclonal antibody and polyclonal serum neutralization assays, as well as receptor- and protease-inhibition tests, suggest that deletion of the FERM-binding motif in the spike CT does not affect antigenicity, receptor affinity, or the overall conformation of the spike protein.

### Application of FERM-binding motif deletion to VLP production

Data in Fig. 3d suggest that truncation of the last 13 or 17 amino acids of the spike CT promotes spike incorporation into VSV pseudoviruses. We therefore hypothesized that these truncations would similarly enhance spike incorporation into virus-like particles (VLPs). To validate this, 293T cells were transfected with plasmids encoding the structural proteins nucleocapsid (N) and membrane (M) from SARS-CoV-1 (**Fig. 5a**) or SARS-CoV-2 (**Fig. 5b**), respectively, together with plasmids expressing spike proteins from SARS-CoV-1, Rs7327, D614G, and GD-pangolin, to generate VLPs. VLPs harvested from the transfected 293T cell supernatants were analyzed via western blot to determine the expression levels of spike, N, and M. Spike incorporation into VLPs was normalized using S/N and S/M ratios, with the FL value set to 1.0. Compared to the full-length spikes, the Δ13 and Δ17 variants of all four viruses showed higher levels of incorporation, with increases ranging from 1.3- to 4.0-fold in SARS-CoV-1 VLPs and 1.1- to 2.9-fold in SARS-CoV-2 VLPs. Notably, for D614G, the Δ13 and Δ17 spikes also exhibited higher cleavage efficiency, leading to increased S2 levels on the VLPs. These results indicate that CT truncation is a viable strategy for enhancing the quality and yield of sarbecovirus VLPs.

**Fig. 5.**
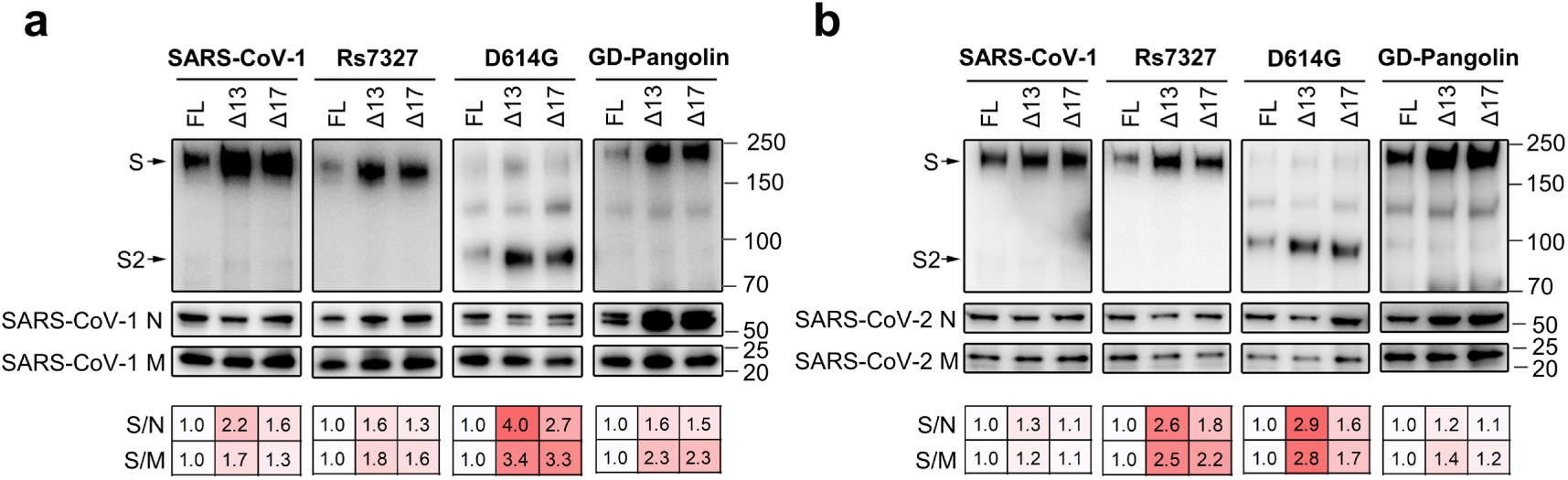
Incorporation of FL, Δ13, and Δ17 spikes into virus-like particles (VLPs). 293T cells were stably transduced with plasmids encoding the structural proteins nucleocapsid (N), membrane (M), and envelope (E) of SARS-CoV-1 (**a**) or SARS-CoV-2 (**b**), followed by transient transfection with plasmids encoding the indicated spike proteins from SARS-CoV-1, Rs7327, SARS-CoV-2 D614G, and GD-pangolin. VLPs were harvested and analyzed by western blot using an anti-S2 antibody cocktail (S2P6, CV3-25, and CC40.8) and an anti-Flag antibody (N and M were C-terminally Flag-tagged). Spike incorporation into VLPs was quantified using ImageJ by measuring band intensities and calculating S/N and S/M ratios. Ratios shown below each band were normalized to FL. Data are representative of two independent experiments.

## Discussion

This work is grounded in a phylogenetically informed paradigm encompassing a broad range of sarbecovirus lineages with defined receptor usage^19, 20, 21^. Representative spike proteins from Clades 1, 3, and 4 were selected for cross-lineage comparative evaluation to dissect how COPI-, COPII-, FERM-, and SNX27-binding motifs within the CT regulate viral infectivity and particle stability. Whereas prior studies have mainly focused on a limited number of viruses, particularly SARS-CoV-1 and SARS-CoV-2, our analysis offers a broader evolutionary perspective on the spike CT and its contribution to viral adaptation^22, 23, 24, 25, 26, 27, 28, 69, 70^. In addition, we examined the impact of CT truncation on viral phenotypes, including sensitivity to neutralization by monoclonal antibodies and sera, as well as susceptibility to small-molecule antivirals (**Fig. 4**). We further assessed the utility of motif-based CT modifications in VLP systems (**Fig. 5**). In summary, our work expands the understanding of sarbecovirus spike CT function, providing practical tools and a theoretical framework for evaluating broadly neutralizing antibodies and small-molecule antivirals, and enabling the proactive design of next-generation vaccines.

To date, the molecular mechanisms by which CT modifications modulate pseudovirus titers remain incompletely understood. Moderate shortening of the CT is widely thought to boost pseudovirus infectivity by yielding a higher density of spike proteins on the virion surface. Two explanations have been put forward. First, deletion of the COPI-binding motif is believed to facilitate spike trafficking to the plasma membrane, thereby elevating its surface abundance. However, multiple studies have shown that mutation or deletion of the COPI binding motif alone has no marked effect on pseudovirus titers^48, 71, 72, 73^. Second, it has been suggested that CT truncation does not substantially alter surface spike expression but instead drives infectivity by reducing S1 subunit shedding, thereby increasing the proportion of functional spike proteins incorporated into virions and thereby strengthening infectivity^46, 47, 72, 73^. Building on these proposed models, we systematically examined spike distribution and processing across CT-shortened variants at three levels: intracellular compartments, the cell surface, and viral particles (**Fig. 3**), and found that removal of the FERM-binding motif alone consistently elevated spike content and cleavage efficiency across these compartments relative to the FL spike, resulting in higher pseudovirus titers and improved particle stability (**Figs. 1–3**). In contrast, truncation of COPI- or SNX27-related motifs was generally detrimental to spike expression, processing, or viral particle stability, whereas truncation of the COPII-binding motif had slightly appreciable impact (**Fig. 1**). Based on these findings, we postulate that the FERM-binding motif mediates interactions between spike and ERM (ezrin/radixin/moesin) proteins, anchoring spike to the actin cytoskeleton and restricting its lateral mobility within the membrane^40^; Disruption of this motif alleviates these constraints, thereby enabling redistribution to viral budding sites and promoting incorporation into virions, ultimately improving pseudovirus infectivity. In conclusion, the FERM-binding motif represents a key limiting element within the CT for pseudovirus production, and its removal provides a molecular basis for augmented pseudovirus titers.

While truncation of the CT is associated with a pronounced augmentation in pseudovirus titers, a central concern for the widespread implementation of this approach is whether it adversely affects the functional integrity of the spike ectodomain. Accumulating evidence suggests that such modifications exert minimal influence on antigenic properties. For instance, Chen et al. reported that truncation of the CT does not alter sike sensitivity to convalescent sera^47^. Yu et al. further demonstrated that full-length SARS-CoV-2 spike and its truncated counterpart display comparable responsiveness to purified antibodies from convalescent sera, soluble human ACE2, and the endosomal inhibitor Bafilomycin A1 (BafA1)^72^. Moreover, Li et al. showed that point mutations within the FERM-binding motif in the SARS-CoV-2 Omicron BA.1 variant increase spike density on VLPs and enhance immunogenicity without affecting neutralization sensitivity^27^. Cumulatively, these observations support the notion that the CT primarily participates in intracellular trafficking and virion assembly, with negligible influence on the antigenicity mediated by the ectodomain. Building on this foundation, we conducted a systematic assessment of how CT shortening influences ectodomain-associated properties. Using serum and antibody neutralization assays, soluble ACE2 inhibition analyses, and small-molecule antiviral assays, we consistently found that the Δ13 and Δ17 variants with FERM truncation enhance pseudovirus titers while exhibiting no detectable differences from the FL construct in antigenicity or receptor-binding capacity (**Fig. 4**). Notably, extension of this strategy to VLP systems revealed that Δ13/Δ17 variants markedly improve spike incorporation into SARS-CoV and SARS-CoV-2 VLPs (**Fig. 5**). These results are collectively summarized in **Fig. 6**. In light of prior evidence linking antigen density to immunogenicity, CT truncation may potentiate immune responses and provide a rationale for vaccine optimization.

**Fig. 6.**
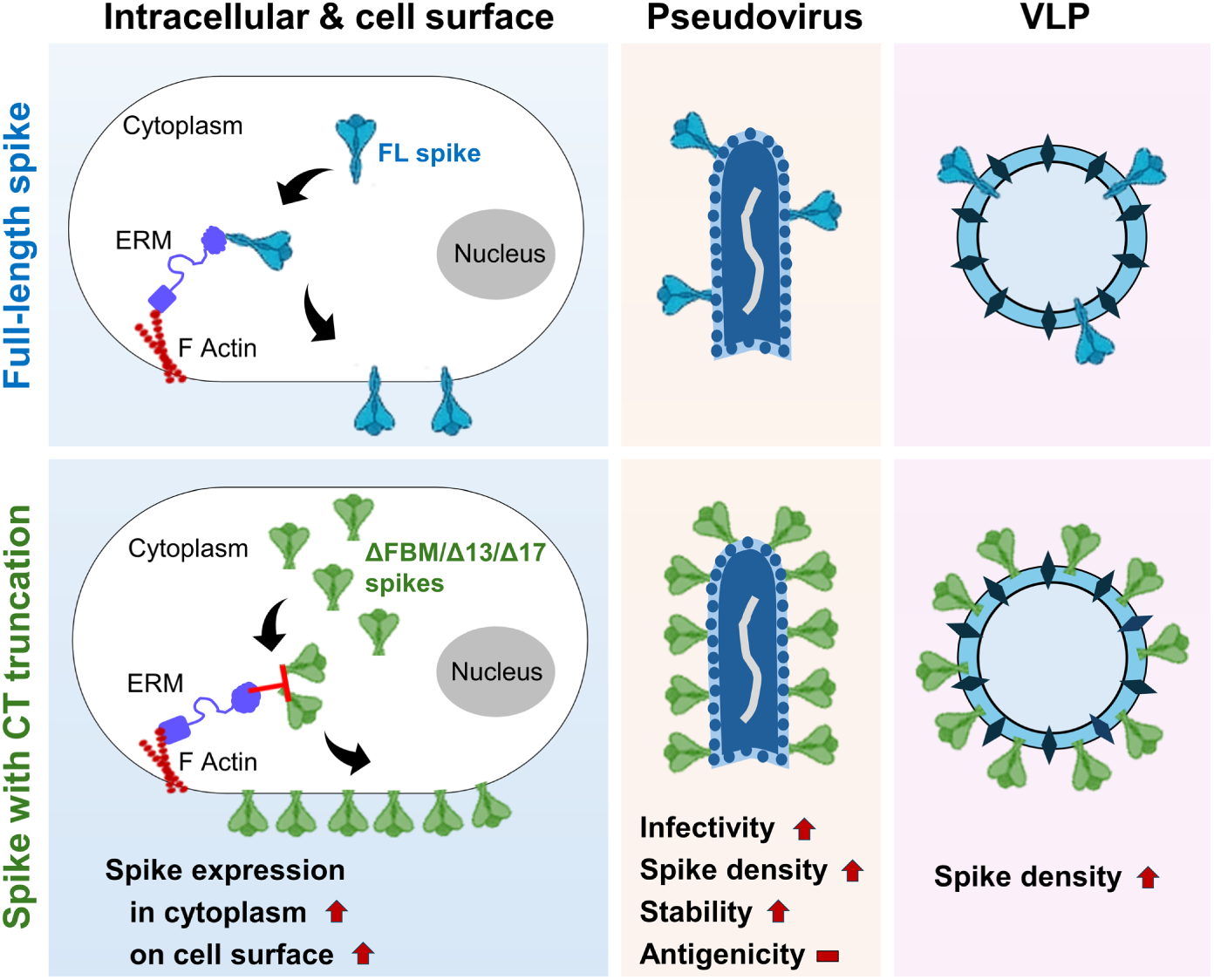
Summary of the effects of CT truncation on spike expression, VSV pseudotyped virus properties, and VLP incorporation. The CT of the sarbecovirus spike protein harbors a FERM-binding motif (FBM), which interacts with the N-terminal domain of the ERM protein family, including ezrin, radixin, and moesin. The C-terminus of the ERM proteins subsequently anchors the spike protein to the cell membrane by interacting with F-actin (filamentous actin). Truncation mutations of the spike protein’s CT (ΔFBM, Δ13, and Δ17) disrupt its interaction with ERM family members, such as moesin. Our findings demonstrate that these CT truncations significantly increase spike expression both intracellularly and on the cell surface. Furthermore, these CT truncations markedly enhance the infectivity of pseudoviruses, increase the spike density on the pseudoviral particles, and improve their thermostability across various temperatures, without altering their antigenicity. Beyond pseudotyped systems, Spike CT truncations also facilitate incorporation into SARS-CoV-1 and SARS-CoV-2 VLPs, thereby increasing the spike density.

Overall, this study delineates the roles of CT motifs in the sarbecovirus spike protein in regulating viral assembly and infectivity. In particular, the FERM-binding motif is identified as a central constraint on these processes, supporting a model in which cytoskeletal interactions restrict spike redistribution. Relief of this limiting factor via CT truncation enhances the assembly efficiency of functional spike proteins in both pseudovirus and VLP systems, without compromising antigenicity or receptor engagement. Our findings establish a mechanistic and translational foundation for improving pseudovirus-based neutralization platforms and guiding vaccine optimization through antigen density modulation, with implications for emerging viral threats and pandemic preparedness.

## Materials and Methods

### Clinical cohort

Seven serum samples were collected in Wuhan, China, between May and October 2025 during the post-COVID-19 pandemic period from individuals without known respiratory infections. Written informed consent was obtained from all participants in accordance with protocols approved by the Hubei Provincial Center for Disease Control and Prevention (Approval No.: HBCDC-AF/SC-08/02.0). Clinical characteristics are summarized in **Table S1**.

### Cell lines

HEK293T (Cat# CRL-3216), Vero E6 (Cat# CRL-1586), and I1-Hybridoma (Cat# CRL-2700) cells were purchased from the American Type Culture Collection (ATCC). Expi293F (Cat# GDC0655) cells were obtained from the China Center for Type Culture Collection (CCTCC). 293T-ACE2, 293T-ACE2-TMPRSS2, Vero E6-ACE2, Vero E6-TMPRSS2, Vero E6-ACE2-TMPRSS2, 293T-SARS-CoV-1-M/N/E, and 293T-SARS-CoV-2-M/N/E cell lines were generated by lentiviral-mediated gene transduction. All adherent cells were cultured in Dulbecco’s modified Eagle medium (DMEM, Gibco, Cat# C11995500BT) containing 10% fetal bovine serum (FBS, HyClone, Cat# SH30406.05) and 100 μg/mL penicillin-streptomycin (Gibco, Cat# 15140122) at 37 °C with 5% CO₂. Expi293F cells were maintained in suspension culture using SMM 293-TII Expression Medium (Sino Biological, Cat# M293TII) supplemented with penicillin-streptomycin at 37 °C under 5% CO₂, with constant shaking at 125 rpm. I1-Hybridoma cells were cultured in Minimum Essential Medium (MEM, Gibco, Cat# C11095500BT) supplemented with 20% fetal bovine serum and 100 μg/mL penicillin-streptomycin at 37 °C with 5% CO₂.

### Plasmids

Genes encoding the spike (S) glycoproteins of SARS-CoV-1, Rs4231, Rs4084, SHC014, Rs7327, WIV1, SARS-CoV-2, RaTG13, GX-Pangolin, GD-Pangolin, BM48-31, Khosta2, and RmYN05 were kindly provided by Dr. Huan Yan at Wuhan University, Wuhan, China, as well as by Dr. Qihui Wang and Dr. George F. Gao at the CAS Key Laboratory of Pathogen Microbiology and Immunology, Institute of Microbiology, Chinese Academy of Sciences (CAS), Beijing, China, and were subsequently cloned into a pCMV vector. Spike protein mutants with deletions of 5,11,13,17 amino acids (Δ5, Δ11, Δ13, Δ17) or FERM-binding motif (ΔFERM) at the C-terminus were generated using a site-directed mutagenesis kit (Vazyme, Cat# C214-02). Additionally, plasmids expressing the SARS-CoV-1 or SARS-CoV-2 M, N, and E proteins with a C-terminal Flag tag, along with constructs for ACE2 and TMPRSS2, were obtained from Professor Pei-Hui Wang at Shandong University, Jinan, China, and were subsequently cloned into a pLVX-Puro vector.

### Generation of stable cell lines

HEK293T cells were seeded into 10 cm dishes 24 h prior to transfection. When cells reached approximately 80% confluence, they were co-transfected with 20 μg of target protein plasmids cloned into pLVX-Puro vectors, along with 15 μg of psPAX2 (Addgene; Cat# 12260) and 5 μg of pMD2.G (Addgene; Cat# 12259), using polyethylenimine (PEI, Polyscience, Cat# 23966-1) as the transfection reagent. After 24 hours of plasmid transfection, the medium was replaced with fresh medium. Cells were cultured for an additional 48 hours, after which the cell culture supernatant containing the recombinant virus was collected. To purify the viral particles, the supernatant was first centrifuged at 600 × *g* for 10 minutes to remove cell debris. Subsequently, it was subjected to ultracentrifugation at 52,000 × *g* to pellet the lentiviral particles. Most of the supernatant was then removed, reserving 1 mL to resuspend the particles. Target cells were then infected with lentivirus-containing suspension. Following integration of the exogenous genome, cells were expanded in culture and positive cells were selected via flow cytometry. The screened cells were required to undergo flow cytometry validation after expansion to ensure suitability for subsequent experiments.

### Production of VSV pseudotyped viruses

HEK293T cells were seeded into 6-well plates and grown overnight, then, 3 μg of spike protein expression plasmid was transfected into HEK293T cells using PEI when the cells reached approximately 80% confluency. After overnight incubation at 37 °C under 5% CO₂, cells were infected with pseudotyped ΔG-luciferase rVSV (Kerafast, Cat# EH1020-PM) at a multiplicity of infection (MOI) of 3 for 2 h, followed by three washes with DMEM containing 2% FBS. Fresh complete culture medium was then added, and cells were incubated for an additional 24 hours at 37 °C. The supernatants containing pseudoviruses were collected and clarified by centrifugation at 600 × *g* for 10 minutes to remove cell debris. Subsequently, I1-hybridoma cell culture supernatant (15% v/v), containing the anti-VSV-G antibody, was added to neutralize residual VSV-G-pseudotyped ΔG-luciferase virus contamination. Pseudotyped viruses were then aliquoted and stored at −80 °C until use.

### Production of lentiviral pseudotyped viruses

For lentiviral pseudovirus production, HEK293T cells were seeded into 6-well plates and cultured overnight as described above. The cells were then co-transfected with 1.5 μg of spike protein expression plasmid and 1.5 μg of HIV-1 backbone luciferase reporter vector (pHIV-1_NL4-3_ ΔEnv-Luc) using PEI. The supernatants were harvested 48 hours post-transfection and centrifuged at 600 × *g* for 10 minutes to remove cell debris. The resulting pseudovirus-containing supernatants were aliquoted and stored at −80 °C for later use.

### Pseudovirus infectivity assay

Pseudovirus titers were assessed by measuring luciferase activity in the target cells. In detail, the virus was 3-fold serially diluted and then incubated with the target cells in a cell culture incubator at 37 °C and 5% CO2. Following 16-24 hours of incubation, luciferase activity was detected using a luciferase assay kit (Promega, Cat# E4550) on a Varioskan LUX microplate reader (Thermo Fisher Scientific).

### Cell surface expression of human ACE2 and TMPRSS2

Cells were digested with trypsin and washed once with FACS buffer (PBS containing 2% FBS). A total of 5×10^5^ cells were resuspended in 100 μL FACS buffer and immediately incubated with anti-human ACE2 antibody (dilution 1:500; Abcam, Cat# ab272500) or anti-human TMPRSS2 antibody (dilution 1:200; Abcam, Cat# ab280567) at 4 °C for 30–45 minutes. Then, cells were incubated with 100 μL of diluted iFluor^TM^ 488 Conjugated Goat anti-rabbit IgG secondary antibody (dilution 1:1,000; HUABIO, Cat# HA1121) in the dark on ice for 30-45 minutes. Subsequently, the cells were resuspended in 200 μL FACS buffer. Cells were washed three times with FACS buffer between steps. ACE2 and TMPRSS2 expression levels were determined using a flow cytometer (Beckman Coulter, CA, USA) and analyzed using FlowJo software v10.9.

### Protein expression

To produce monoclonal antibodies VIR-7229^62^, 10-40^60^, SA55^63^, VYD222^64, 65^, S309^61^, S2P6^57^, CV3-25^58^, and CC40.8^59^, Expi293F cells were co-transfected with the corresponding light chain and heavy chain expression plasmids at a 1:1 ratio using PEI. Five days post-transfection, cell supernatants were harvested and antibodies were purified by affinity chromatography using Protein A resin (GenScript, Cat# L00210-50).

To produce the soluble human ACE2 protein fused to an Fc tag (hACE2-Fc), pcDNA3-sACE2-WT(732)-IgG1 plasmid (Addgene; Cat# 154104) was transfected into Expi293F cells. hACE2-Fc was purified using Protein A resin as well.

### Western blot

For cell lysates, HEK293T cells were transfected with the S glycoprotein-expressing plasmids for 48 hours. The cells were then washed with 1× PBS buffer and lysed with 1% NP-40 buffer (Sangon Biotech, CAS# 9016-45-9) on ice for 10 minutes. Lysates were centrifuged at 16,100 × *g* for 10 minutes at 4 °C, and supernatants were mixed with 5× loading buffer followed by boiling at 95 °C for 10 minutes.

For pseudotyped virus, VSV-based pseudoviruses were prepared as described above. Cell supernatants containing pseudoviruses were collected, clarified by low-speed centrifugation (600 × *g* for 10 minutes), and then centrifuged at 16,100 × *g* for 1 hour at 4 °C. After centrifugation, supernatants were aspirated, and viral pellets were resuspended and boiled in 1× loading buffer at 95 °C for 10 minutes.

Protein samples from cell lysates and pseudovirus pellets were subjected to sodium dodecyl sulfate–polyacrylamide gel electrophoresis (SDS-PAGE) and western blotting. Spike proteins were detected using a cocktail of monoclonal antibodies S2P6^57^, CV3-25^58^, and CC40.8^59^. HRP-conjugated goat anti-human IgG1 secondary antibody (ABclonal, Cat# AS002) was used for chemiluminescent detection. GAPDH and VSV-N proteins were detected using GAPDH mouse mAb (ABclonal, Cat# AC002) and Anti-VSV-N mouse mAb (Absolute Antibody, Ab01403-2.0) as primary antibodies respectively, followed by HRP-conjugated goat anti-mouse IgG secondary antibody (ABclonal, Cat# AS003). Flag-tagged M, N, and E proteins were detected using DYKDDDDK Tag (9A3) Mouse mAb (Cell Signaling Technology, Cat# 8146T) as the primary antibody, and HRP-conjugated Goat anti-Mouse IgG (ABclonal, Cat# AS003) as the secondary antibody.

### Cell surface expression of spike protein

The expression of spike protein on the cell surface was quantified by flow cytometry. Plasmids expressing the spike proteins of SARS-CoV-1, Rs7327, D614G and GD-Pangolin along with their variants featuring CT truncations, were transfected into Expi293F cells at a 1:1 ratio with the pRRLSIN.cPPT.PGK-GFP.WPRE plasmid using PEI. At 48 hours post-transfection, cells were harvested, resuspended in FACS buffer to a concentration of 5×10^6^ cells/mL, and then incubated with serial dilutions of the test antibodies or hACE2-Fc (starting from an initial concentration of 10 μg/mL, in 4-fold serial dilutions) at 4 °C for 30–45 minutes. Following this, the cells were washed three times with FACS buffer and incubated with APC anti-human IgG Fc secondary antibody (Biolegend, Cat# 410712) for 30-45 minutes at 4 °C. Finally, the cells were washed three times with FACS buffer, resuspended, and analyzed using a flow cytometer.

### Thermal stability and infectivity of spike variants

Freshly prepared pseudoviruses were incubated in an ice-water bath (0 °C), or in water baths maintained at 25 °C and 37 °C, for the indicated durations (0, 12, 24, 48, and 72 h). After incubation, the pseudoviruses were titrated on Vero E6-ACE2-TMPRSS2 cells according to the aforementioned pseudovirus infectivity assay method.

### Neutralization assays

VSV-based pseudoviruses bearing either wild-type or CT truncated spike proteins from SARS-CoV-1, Rs7327, D614G, or GD-Pangolin were produced as described above. To determine their sensitivity to antibodies and sera, antibodies were diluted in a 5-fold gradient from a working concentration of 10 μg/mL in a 96-well plate, while serum samples were diluted in a 4-fold gradient, starting at a 1:25 dilution. Subsequently, pseudovirus was added and the mixture was incubated at 37 °C for 1 hour. Following incubation, target cells were added and incubated overnight before luciferase activity was measured. Vero E6 cells (4×10⁴ cells/well) were used for SARS-CoV-1, D614G, and GD-Pangolin pseudoviruses, whereas 293T-ACE2 cells (1×10^5^ cells/well) were used for Rs7327 pseudovirus. Neutralization curves and half-maximal inhibitory concentration (IC₅₀) or half-maximal inhibitory dilution (ID₅₀) values were calculated by fitting a nonlinear five-parameter dose-response curve using GraphPad Prism v10.

### Soluble ACE2 inhibition assay

Pseudoviruses were pre-titrated as previously described. Pseudoviruses were incubated with 2-fold serially diluted hACE2-Fc protein (starting from 10 μg/mL) in 96-well plates at 37 °C for 1 hour. Following incubation, target cells were added and incubated overnight before luciferase activity was measured. Vero E6 cells (4×10⁴ cells/well) were used for SARS-CoV-1, D614G, and GD-Pangolin pseudoviruses; 293T-ACE2 cells (1×10^5^ cells/well) were used for Rs7327 pseudovirus. Neutralization curves and half-maximal inhibitory concentration (IC₅₀) values were calculated by fitting a nonlinear five-parameter dose-response curve using GraphPad Prism v10.

### Inhibition of pseudovirus variants by host protease inhibitors

To test the effect of host protease inhibitors on pseudovirus entry, Vero E6-ACE2-TMPRSS2 cells (4×10⁴ cells/well) were pre-incubated for 2 hours with serially diluted cathepsin L inhibitor E64d (MedChemExpress, CAS# 88321-09-9) or TMPRSS2 inhibitor camostat mesylate (MedChemExpress, CAS# 59721-29-8). Pseudoviruses were then added, and luciferase activity was measured after 16-24 hours of incubation. Inhibition curves and half-maximal effective concentration (EC₅₀) values were generated using a nonlinear five-parameter dose-response model in GraphPad Prism v10.

### Virus-like particle assay

To generate virus-like particles (VLPs), HEK293T cells stably expressing Flag-tagged nucleocapsid (N), membrane (M), and envelope (E) proteins of SARS-CoV-1 or SARS-CoV-2 (293T-SARS1-M/N/E and 293T-SARS2-M/N/E) were transiently transfected with spike protein-expressing plasmids. At 48 hours post-transfection, cell culture supernatants were collected and centrifuged at 600 × *g* for 10 minutes at 4 °C to remove cell debris. The clarified supernatants were loaded onto a 20% (v/v) sucrose cushion and centrifuged at 21,000 × *g* for 2 hours at 4 °C. After discarding the supernatants, the pellets were resuspended in 40 μL of 1× PBS buffer. The samples were mixed with 10 μL of 5× protein loading buffer, boiled at 95 °C for 10 minutes, and then analyzed by western blotting.

### Phylogenetic and sequence analysis

Full-length spike glycoprotein sequences of the *Sarbecovirus* subgenus were sourced from the GenBank database. The sequences were aligned with MUSCLE (v5.1)^74^, trimmed by trimAl (v1.4)^75^, and used to infer maximum likelihood phylogenetic trees in IQ-TREE (v2.2.0)^76^. The best-fit substitution model, JTTDCMut+F+I+R3, was selected by ModelFinder^77^ based on the Bayesian Information Criterion, and branch support was assessed with 1,000 ultrafast bootstrap replicates. Final trees were visualized and annotated in the Interactive Tree Of Life (iTOL) v6^78^.

### Co-immunoprecipitation (Co-IP)

For co-immunoprecipitation, plasmids encoding wild-type or mutated D614G spike and Moesin with a C-terminal 2× Strep tag were transfected into HEK293T cells with PEI MAX (1mg/mL) reagent. After 48 h, cells were harvested using 1× PBS buffer and then lysed in RIPA buffer (50 mM Tris-HCl, 150 mM NaCl, 1% NP-40, 0.25% sodium deoxycholate) supplemented with 1× Protease Inhibitor Cocktail (MCE, HY-K0010). A 10% (v/v) aliquot of each lysate was mixed with 5× SDS-PAGE loading buffer (containing 100 mM DTT), boiled at 95°C for 10 min and then used as input control. The remaining lysates were incubated with 5 μg of anti-D614G spike mAb (SA55, in-house), under gentle rotation at 4℃ for 1 h in 1.5 mL microcentrifuge tubes. Then, 25 μL Protein A/G magnetic beads (MCE, HY-K0202) were added into the mixture, and rotated at 4°C overnight. The next day, the beads were washed with 1 mL RIPA buffer three times at 4°C (each wash with 5 min rotation), and then boiled with 30 μL 1× protein loading buffer at 95°C for 10 min. The eluted samples and input controls were then subjected to Western blot.

### Data analysis

IC₅₀, ID₅₀, and EC₅₀ values were determined by fitting the data to five-parameter dose-response curves using GraphPad Prism (v.10). The same software was also used for data visualization.

### Data availability

All experimental data are provided within this manuscript. Sarbecovirus spike sequences were obtained from the National Center for Biotechnology Information (NCBI) and the Global Initiative on Sharing All Influenza Data (GISAID) databases (https://www.gisaid.org/). All data supporting the findings of this study are available in the article, the Supplementary Information, and the Source Data files. Source Data are provided with this paper.

## Acknowledgements

This work was supported by the National Natural Science Foundation of China (grant no. 92569102 to L.L.). Additional support was provided by the Taikang Center for Life and Medical Sciences, Wuhan University, Wuhan, China, and the State Key Laboratory of Virology and Biosafety, Wuhan University, Wuhan, China, for L.L. and Q.W. The funders had no role in study design, data collection and analysis, decision to publish, or preparation of the manuscript.

## Author Contributions

Q.W. and L.L. conceived and supervised the project. S.Z., C.C., L.S. and X.Z. performed the experiments. L.M. and P.C. helped collect clinical samples. C.L. conducted bioinformatic analyses. Q.W., L.L., S.Z., C.C. and C.L. analyzed the data and wrote the manuscript.

---

**Table S1.** Summary of clinical cohort.

| Sample ID | Vaccine history |  |  |  |  |  | Date of vaccination |  |  |  |  |  | Age | Gender |
| --- | --- | --- | --- | --- | --- | --- | --- | --- | --- | --- | --- | --- | --- | --- |
|  | 1<sup>st</sup> | 2<sup>nd</sup> | 3<sup>rd</sup> | 4<sup>th</sup> | 5<sup>th</sup> | 6<sup>th</sup> | 1<sup>st</sup> | 2<sup>nd</sup> | 3<sup>rd</sup> | 4<sup>th</sup> | 5<sup>th</sup> | 6<sup>th</sup> |  |  |
| 1 | BIBP-CorV | CIBP-CorV | BIBP-CorV | - | - | - | 08.05.2021 | 08.29.2021 | 06.14.2022 | - | - | - | 21 | Female |
| 2 | BIBP-CorV | BIBP-CorV | BIBP-CorV | - | - | - | 08.10.2021 | 08.24.2021 | 08.06.2022 | - | - | - | 21 | Male |
| 3 | CoronaVac | CoronaVac | CoronaVac | - | - | - | 07.16.2021 | 08.19.2021 | 02.19.2022 | - | - | - | 21 | Male |
| 4 | WIBP-CorV | BIBP-CorV | WIBP-CorV | - | - | - | 03.19.2021 | 05.10.2021 | 01.14.2022 | - | - | - | 28 | Male |
| 5 | BIBP-CorV | BIBP-CorV | CIBP-CorV | - | - | - | 08.06.2021 | 08.18.2021 | 07.24.2022 | - | - | - | 21 | Female |
| 6 | BIBP-CorV | BIBP-CorV | BIBP-CorV | - | - | - | 04.24.2021 | 06.11.2021 | 01.06.2022 | - | - | - | 23 | Female |
| 7 | BNT162b2 | BNT162b2 | BNT162b2 | mRNA-1273.222 | mRNA-1273.815 | BNT162b2 | 01.03.2021 | 01.28.2021 | 01.08.2022 | 09.19.2022 | 04.28.2023 | 10.02.2024 | 43 | Male |

**Fig. S1.**
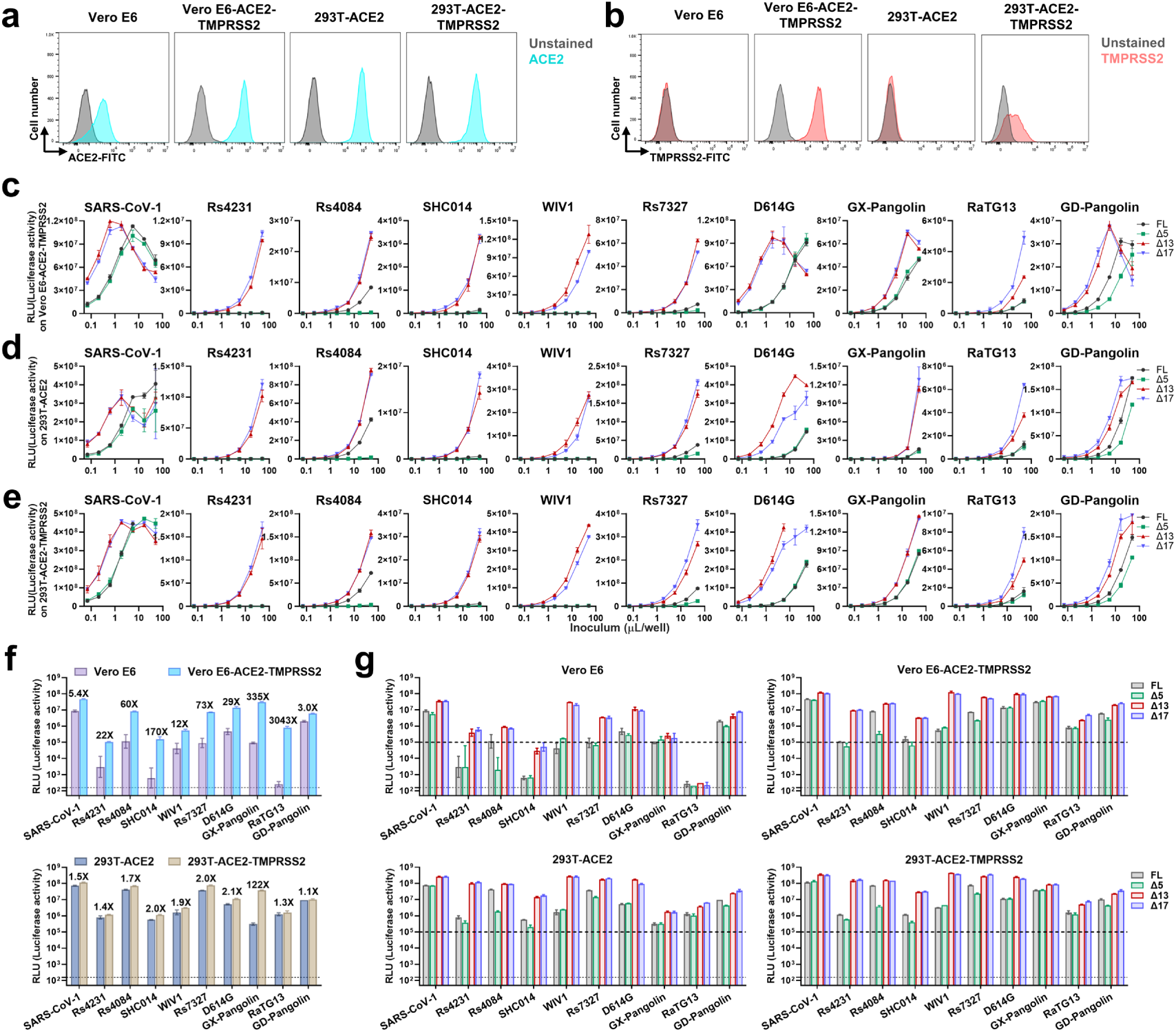
Truncation of the FERM-binding motif increases viral titers, whereas truncation of COPI- and COPII-binding motifs does not. **a.** Cell surface expression of human ACE2 in Vero E6, Vero E6-ACE2-TMPRSS2, 293T-ACE2, and 293T-ACE2-TMPRSS2 cells. **b.** Cell surface expression of human TMPRSS2 in the same cell lines as in a. **c.** Titration of VSV pseudoviruses bearing FL and CT-truncated spike glycoproteins in Vero E6-ACE2-TMPRSS2 cells. Infectivity was assessed by measuring luciferase activity (relative light units, RLU). Data are presented as mean ± s.d. of triplicates. **d.** Titration of VSV pseudoviruses bearing FL and CT-truncated spike glycoproteins in 293T-ACE2 cells **e.** Titration of VSV pseudoviruses bearing FL and CT-truncated spike glycoproteins in 293T-ACE2-TMPRSS2 cells. **f.** Infectivity of VSV pseudoviruses bearing the indicated FL spike proteins in Vero E6, Vero E6-ACE2-TMPRSS2, 293T-ACE2, and 293T-ACE2-TMPRSS2 cells. Fold increases in infectivity in Vero E6-ACE2-TMPRSS2 versus Vero E6 and in 293T-ACE2-TMPRSS2 versus 293T-ACE2 cells are shown. Dashed lines indicate background signal in the absence of virus input. **g.** Infectivity of VSV pseudoviruses bearing FL and CT-truncated spike proteins in Vero E6, Vero E6-ACE2-TMPRSS2, 293T-ACE2, and 293T-ACE2-TMPRSS2 cells. Dashed lines indicate background signal in the absence of virus input.

**Fig. S2.**
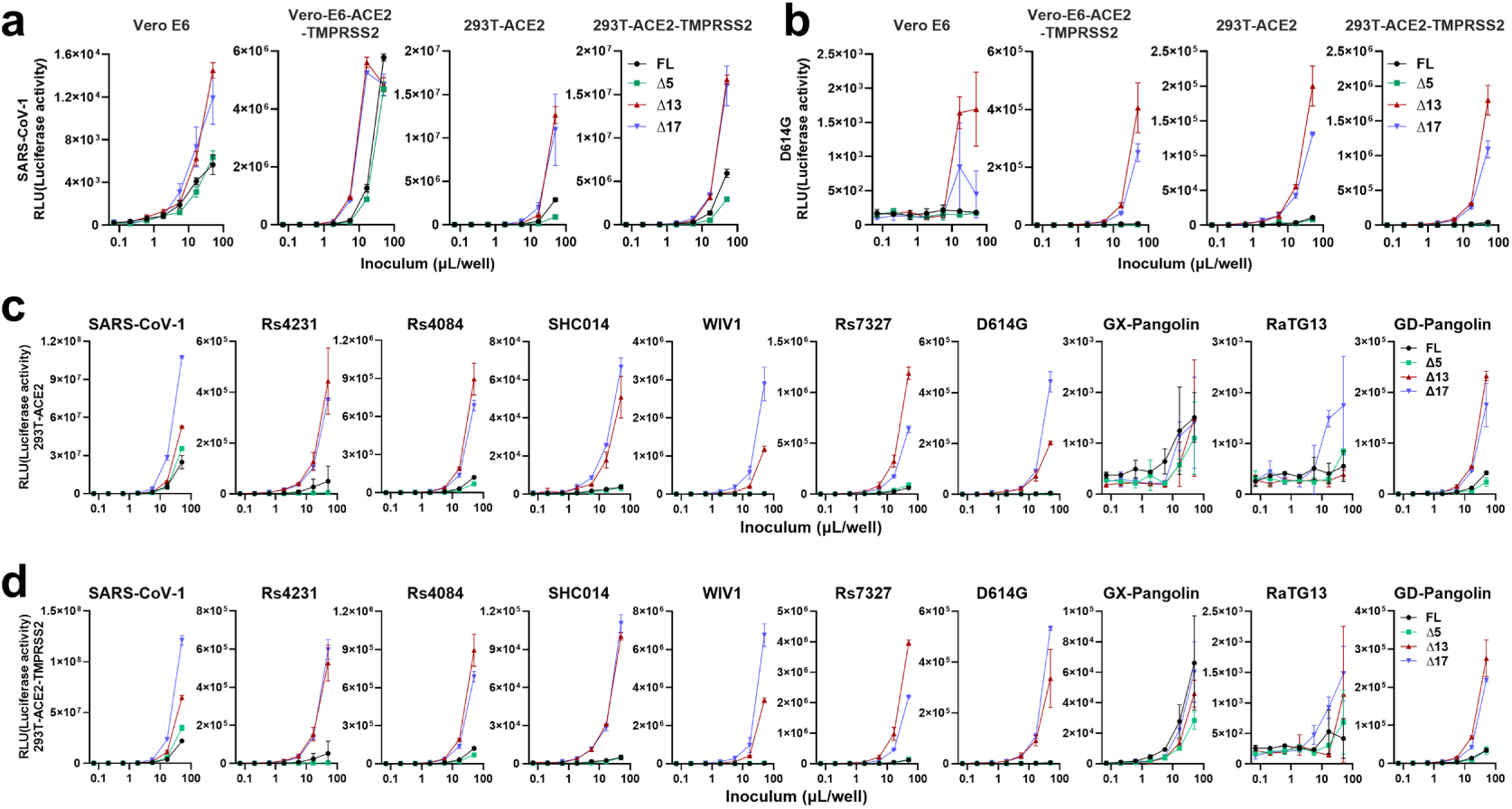
CT truncation of spike increases lentiviral pseudovirus titers of clade 1 sarbecoviruses. **a.** Infectivity of lentivirus pseudotyped with FL or CT-truncated SARS-CoV-1 spike in Vero E6 cells. **b.** Infectivity of lentivirus pseudotyped with FL or CT-truncated SARS-CoV-2 D614G spike in Vero E6 cells. **c.** Infectivity of lentivirus pseudotyped with FL or CT-truncated spikes in 293T-ACE2 cells. **d.** Infectivity of lentivirus pseudotyped with FL or CT-truncated indicated spikes in 293T-ACE2-TMPRSS2 cells.

**Fig. S3.**
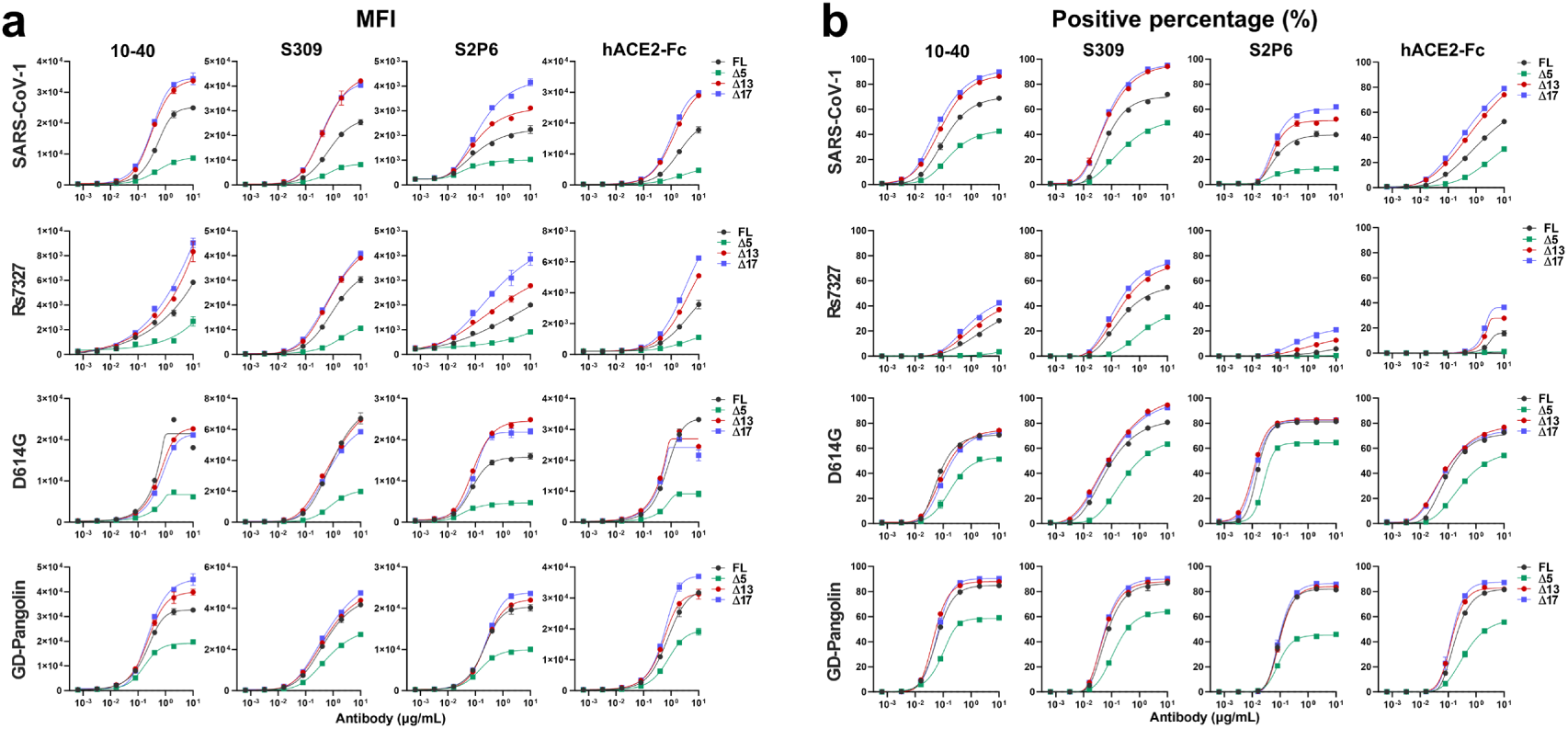
Expression of FL, Δ5, Δ13, and Δ17 spike glycoproteins on Expi293F cells. Expi293F cells were transiently transfected with plasmids encoding the indicated spike glycoproteins and stained with serial dilutions of antibodies or human ACE2 fused to an Fc tag (hACE2-Fc). Mean fluorescence intensity (MFI) (**a**) and percentage of positive cells (**b**) are shown. Data are based on two technical replicates and are representative of two independent experiments, presented as mean ± s.d.

## References

1. Ge XY, et al. Isolation and characterization of a bat SARS-like coronavirus that uses the ACE2 receptor. Nature 503, 535–538 (2013).

2. Zhou P, et al. A pneumonia outbreak associated with a new coronavirus of probable bat origin. Nature 579, 270–273 (2020).

3. Hu B, Guo H, Zhou P, Shi ZL. Characteristics of SARS-CoV-2 and COVID-19. Nat Rev Microbiol 19, 141–154 (2021).

4. Li W, et al. Bats are natural reservoirs of SARS-like coronaviruses. Science 310, 676–679 (2005).

5. Lau SK, et al. Severe acute respiratory syndrome coronavirus-like virus in Chinese horseshoe bats. Proc Natl Acad Sci U S A 102, 14040–14045 (2005).

6. Guan Y, et al. Isolation and characterization of viruses related to the SARS coronavirus from animals in southern China. Science 302, 276–278 (2003).

7. Ksiazek TG, et al. A novel coronavirus associated with severe acute respiratory syndrome. N Engl J Med 348, 1953–1966 (2003).

8. Zhu N, et al. A Novel Coronavirus from Patients with Pneumonia in China, 2019. N Engl J Med 382, 727–733 (2020).

9. ZLi F. Structure, Function, and Evolution of Coronavirus Spike Proteins. Annu Rev Virol 3, 237–261 (2016).

10. Beniac DR, Andonov A, Grudeski E, Booth TF. Architecture of the SARS coronavirus prefusion spike. Nat Struct Mol Biol 13, 751–752 (2006).

11. Gui M, et al. Cryo-electron microscopy structures of the SARS-CoV spike glycoprotein reveal a prerequisite conformational state for receptor binding. Cell Res 27, 119–129 (2017).

12. Bosch BJ, van der Zee R, de Haan CA, Rottier PJ. The coronavirus spike protein is a class I virus fusion protein: structural and functional characterization of the fusion core complex. J Virol 77, 8801–8811 (2003).

13. Walls AC, Park YJ, Tortorici MA, Wall A, McGuire AT, Veesler D. Structure, Function, and Antigenicity of the SARS-CoV-2 Spike Glycoprotein. Cell 181, 281–292 e286 (2020).

14. Wrapp D, et al. Cryo-EM structure of the 2019-nCoV spike in the prefusion conformation. Science 367, 1260–1263 (2020).

15. Du L, He Y, Zhou Y, Liu S, Zheng BJ, Jiang S. The spike protein of SARS-CoV--a target for vaccine and therapeutic development. Nat Rev Microbiol 7, 226–236 (2009).

16. Salvatori G, et al. SARS-CoV-2 SPIKE PROTEIN: an optimal immunological target for vaccines. J Transl Med 18, 222 (2020).

17. Hutchinson GB, et al. Nanoparticle display of prefusion coronavirus spike elicits S1-focused cross-reactive antibody response against diverse coronavirus subgenera. Nat Commun 14, 6195 (2023).

18. Piccoli L, et al. Mapping Neutralizing and Immunodominant Sites on the SARS-CoV-2 Spike Receptor-Binding Domain by Structure-Guided High-Resolution Serology. Cell 183, 1024–1042 e1021 (2020).

19. Letko M, Marzi A, Munster V. Functional assessment of cell entry and receptor usage for SARS-CoV-2 and other lineage B betacoronaviruses. Nat Microbiol 5, 562–569 (2020).

20. Starr TN, et al. SARS-CoV-2 RBD antibodies that maximize breadth and resistance to escape. Nature 597, 97–102 (2021).

21. Starr TN, et al. ACE2 binding is an ancestral and evolvable trait of sarbecoviruses. Nature 603, 913–918 (2022).

22. Cattin-Ortola J, Welch LG, Maslen SL, Papa G, James LC, Munro S. Sequences in the cytoplasmic tail of SARS-CoV-2 Spike facilitate expression at the cell surface and syncytia formation. Nat Commun 12, 5333 (2021).

23. Dey D, et al. An extended motif in the SARS-CoV-2 spike modulates binding and release of host coatomer in retrograde trafficking. Commun Biol 5, 115 (2022).

24. Wu Z, et al. Peptide targeting the interaction of S protein cysteine-rich domain with Ezrin restricts pan-coronavirus infection. Signal Transduct Target Ther 8, 19 (2023).

25. Yang B, et al. SNX27 suppresses SARS-CoV-2 infection by inhibiting viral lysosome/late endosome entry. Proc Natl Acad Sci U S A 119, (2022).

26. Ren Y, Liu Y, Zhang Z, Liu Y, Li K, Zhang L. SNX27-mediated endocytic recycling of GLUT1 is suppressed by SARS-CoV-2 spike, possibly explaining neuromuscular disorders in patients with COVID-19. J Infect 85, e116–e118 (2022).

27. Li Y, et al. A substitution at the cytoplasmic tail of the spike protein enhances SARS-CoV-2 infectivity and immunogenicity. EBioMedicine 110, 105437 (2024).

28. Wang J, et al. SARS-CoV-2 S assembly into virions facilitated by host ERM proteins. Proc Natl Acad Sci U S A 123, e2504517123 (2026).

29. Gomez-Navarro N, Miller E. Protein sorting at the ER-Golgi interface. J Cell Biol 215, 769–778 (2016).

30. Letourneur F, et al. Coatomer is essential for retrieval of dilysine-tagged proteins to the endoplasmic reticulum. Cell 79, 1199–1207 (1994).

31. Brandizzi F, Barlowe C. Organization of the ER-Golgi interface for membrane traffic control. Nat Rev Mol Cell Biol 14, 382–392 (2013).

32. Jackson LP, et al. Molecular basis for recognition of dilysine trafficking motifs by COPI. Dev Cell 23, 1255–1262 (2012).

33. Ma W, Goldberg J. Rules for the recognition of dilysine retrieval motifs by coatomer. EMBO J 32, 926–937 (2013).

34. Lontok E, Corse E, Machamer CE. Intracellular targeting signals contribute to localization of coronavirus spike proteins near the virus assembly site. J Virol 78, 5913–5922 (2004).

35. McBride CE, Li J, Machamer CE. The cytoplasmic tail of the severe acute respiratory syndrome coronavirus spike protein contains a novel endoplasmic reticulum retrieval signal that binds COPI and promotes interaction with membrane protein. J Virol 81, 2418–2428 (2007).

36. Sadasivan J, Singh M, Sarma JD. Cytoplasmic tail of coronavirus spike protein has intracellular targeting signals. J Biosci 42, 231–244 (2017).

37. Jennings BC, Kornfeld S, Doray B. A weak COPI binding motif in the cytoplasmic tail of SARS-CoV-2 spike glycoprotein is necessary for its cleavage, glycosylation, and localization. FEBS Lett 595, 1758–1767 (2021).

38. Votsmeier C, Gallwitz D. An acidic sequence of a putative yeast Golgi membrane protein binds COPII and facilitates ER export. EMBO J 20, 6742–6750 (2001).

39. Millet JK, et al. Ezrin interacts with the SARS coronavirus Spike protein and restrains infection at the entry stage. PLoS One 7, e49566 (2012).

40. Ponuwei GA. A glimpse of the ERM proteins. J Biomed Sci 23, 35 (2016).

41. Fehon RG, McClatchey AI, Bretscher A. Organizing the cell cortex: the role of ERM proteins. Nat Rev Mol Cell Biol 11, 276–287 (2010).

42. Lunn ML, et al. A unique sorting nexin regulates trafficking of potassium channels via a PDZ domain interaction. Nat Neurosci 10, 1249–1259 (2007).

43. Gallon M, et al. A unique PDZ domain and arrestin-like fold interaction reveals mechanistic details of endocytic recycling by SNX27-retromer. Proc Natl Acad Sci U S A 111, E3604–3613 (2014).

44. Petit CM, et al. Palmitoylation of the cysteine-rich endodomain of the SARS-coronavirus spike glycoprotein is important for spike-mediated cell fusion. Virology 360, 264–274 (2007).

45. Giroglou T, et al. Retroviral vectors pseudotyped with severe acute respiratory syndrome coronavirus S protein. J Virol 78, 9007–9015 (2004).

46. Havranek KE, et al. SARS-CoV-2 Spike Alterations Enhance Pseudoparticle Titers and Replication-Competent VSV-SARS-CoV-2 Virus. Viruses 12, (2020).

47. Chen HY, et al. Cytoplasmic Tail Truncation of SARS-CoV-2 Spike Protein Enhances Titer of Pseudotyped Vectors but Masks the Effect of the D614G Mutation. J Virol 95, e0096621 (2021).

48. Hu L, et al. A new intracellular targeting motif in the cytoplasmic tail of the spike protein may act as a target to inhibit SARS-CoV-2 assembly. Antiviral Res 209, 105509 (2023).

49. Liu P, et al. Design of customized coronavirus receptors. Nature 635, 978–986 (2024).

50. Ogando NS, et al. SARS-coronavirus-2 replication in Vero E6 cells: replication kinetics, rapid adaptation and cytopathology. J Gen Virol 101, 925–940 (2020).

51. Schmidt F, et al. Measuring SARS-CoV-2 neutralizing antibody activity using pseudotyped and chimeric viruses. J Exp Med 217, (2020).

52. Nie J, et al. Establishment and validation of a pseudovirus neutralization assay for SARS-CoV-2. Emerg Microbes Infect 9, 680–686 (2020).

53. Jackson CB, Farzan M, Chen B, Choe H. Mechanisms of SARS-CoV-2 entry into cells. Nat Rev Mol Cell Biol 23, 3–20 (2022).

54. Roelle SM, Shukla N, Pham AT, Bruchez AM, Matreyek KA. Expanded ACE2 dependencies of diverse SARS-like coronavirus receptor binding domains. PLoS Biol 20, e3001738 (2022).

55. Islam A, et al. Spatial epidemiology and genetic diversity of SARS-CoV-2 and related coronaviruses in domestic and wild animals. PLoS One 16, e0260635 (2021).

56. Neerukonda SN, et al. Establishment of a well-characterized SARS-CoV-2 lentiviral pseudovirus neutralization assay using 293T cells with stable expression of ACE2 and TMPRSS2. PLoS One 16, e0248348 (2021).

57. Pinto D, et al. Broad betacoronavirus neutralization by a stem helix-specific human antibody. Science 373, 1109–1116 (2021).

58. Hurlburt NK, et al. Structural definition of a pan-sarbecovirus neutralizing epitope on the spike S2 subunit. Commun Biol 5, 342 (2022).

59. Zhou P, et al. A human antibody reveals a conserved site on beta-coronavirus spike proteins and confers protection against SARS-CoV-2 infection. Sci Transl Med 14, eabi9215 (2022).

60. Liu L, et al. An antibody class with a common CDRH3 motif broadly neutralizes sarbecoviruses. Sci Transl Med 14, eabn6859 (2022).

61. Pinto D, et al. Cross-neutralization of SARS-CoV-2 by a human monoclonal SARS-CoV antibody. Nature 583, 290–295 (2020).

62. Rosen LE, et al. A potent pan-sarbecovirus neutralizing antibody resilient to epitope diversification. Cell 187, 7196–7213.e7126 (2024).

63. Cao Y, et al. Rational identification of potent and broad sarbecovirus-neutralizing antibody cocktails from SARS convalescents. Cell Rep 41, 111845 (2022).

64. Ison MG, et al. Efficacy and Safety of Adintrevimab (ADG20) for the Treatment of High-Risk Ambulatory Patients With Mild or Moderate Coronavirus Disease 2019: Results From a Phase 2/3, Randomized, Placebo-Controlled Trial (STAMP) Conducted During Delta Predominance and Early Emergence of Omicron. Open Forum Infect Dis 10, ofad279 (2023).

65. Yuan M, et al. Structural and functional analysis of VYD222: a broadly neutralizing antibody against SARS-CoV-2 variants. bioRxiv, (2025).

66. Hoffmann M, et al. SARS-CoV-2 Cell Entry Depends on ACE2 and TMPRSS2 and Is Blocked by a Clinically Proven Protease Inhibitor. Cell 181, 271–280.e278 (2020).

67. Sun G, et al. Structural Basis of Covalent Inhibitory Mechanism of TMPRSS2-Related Serine Proteases by Camostat. J Virol 95, e0086121 (2021).

68. Zhao MM, et al. Cathepsin L plays a key role in SARS-CoV-2 infection in humans and humanized mice and is a promising target for new drug development. Signal Transduct Target Ther 6, 134 (2021).

69. Burkova EE, Bakhno IA. Sequences in the Cytoplasmic Tail Contribute to the Intracellular Trafficking and the Cell Surface Localization of SARS-CoV-2 Spike Protein. Biomolecules 15, (2025).

70. Kong W, et al. Disruption of spike protein N-glycosylation induces its endoplasmic reticulum retention and attenuates SARS-CoV-2 infectivity. J Virol 100, e0027026 (2026).

71. Hirabayashi A, et al. Coatomer complex I is required for the transport of SARS-CoV-2 progeny virions from the endoplasmic reticulum-Golgi intermediate compartment. mBio 16, e0333124 (2025).

72. Yu J, et al. Deletion of the SARS-CoV-2 Spike Cytoplasmic Tail Increases Infectivity in Pseudovirus Neutralization Assays. J Virol 95, (2021).

73. Zhang L, et al. Cytoplasmic Tail Truncation Stabilizes S1-S2 Association and Enhances S Protein Incorporation into SARS-CoV-2 Pseudovirions. J Virol 97, e0165022 (2023).

74. Edgar RC. Muscle5: High-accuracy alignment ensembles enable unbiased assessments of sequence homology and phylogeny. Nat Commun 13, 6968 (2022).

75. Capella-Gutierrez S, Silla-Martinez JM, Gabaldon T. trimAl: a tool for automated alignment trimming in large-scale phylogenetic analyses. Bioinformatics 25, 1972–1973 (2009).

76. Minh BQ, et al. Corrigendum to: IQ-TREE 2: New Models and Efficient Methods for Phylogenetic Inference in the Genomic Era. Mol Biol Evol 37, 2461 (2020).

77. Kalyaanamoorthy S, Minh BQ, Wong TKF, von Haeseler A, Jermiin LS. ModelFinder: fast model selection for accurate phylogenetic estimates. Nat Methods 14, 587–589 (2017).

78. Letunic I, Bork P. Interactive Tree of Life (iTOL) v6: recent updates to the phylogenetic tree display and annotation tool. Nucleic Acids Res 52, W78–W82 (2024).

